# Mannose-coupled AAV2: a second generation AAV vector for increased retinal gene therapy efficiency

**DOI:** 10.1101/2022.12.01.518481

**Authors:** Mathieu Mével, Virginie Pichard, Mohammed Bouzelha, Dimitri Alvarez-Dorta, Pierre-Alban Lalys, Nathalie Provost, Marine Allais, Alexandra Mendes, Elodie Landagaray, Jean-Baptiste Ducloyer, Anne Galy, Nicole Brument, Gaëlle M. Lefevre, Sébastien G. Gouin, Carolina Isiegas, Guylène Le Meur, Thérèse Cronin, Caroline Le Guiner, Michel Weber, Philippe Moullier, Eduard Ayuso, David Deniaud, Oumeya Adjali

## Abstract

Inherited retinal diseases are a leading and untreatable cause of blindness and are therefore candidate diseases for gene therapy. Recombinant vectors derived from adeno-associated virus (rAAV) are currently the most promising vehicles for *in vivo* therapeutic gene delivery to the retina. However, there is a need for novel AAV-based vectors with greater efficacy for ophthalmic applications, as underscored by recent reports of dose-related inflammatory responses in clinical trials of rAAV-based ocular gene therapies. Improved therapeutic efficacy of vectors would allow for decreases in the dose delivered, with consequent reductions in immune reactions. Here, we describe the development of new rAAV vectors using bioconjugation chemistry to modify the rAAV capsid, thereby improving the therapeutic index. Covalent coupling of a mannose ligand, *via* the formation of a thiourea bond, to the amino groups of the rAAV capsid significantly increases vector transduction efficiency of both rat and nonhuman primate retinas. These optimized rAAV vectors have important implications for the treatment of a wide range of retinal diseases.

## 1. Introduction

Recombinant viral vectors derived from adeno-associated viruses (rAAVs) constitute the most promising *in vivo* gene transfer platform for gene therapy of various human genetic diseases, including hemophilia, muscular dystrophies, and retinal blindness. Indeed, one of only two rAAV products currently on the market is approved for the treatment of RPE65-related Leber congenital amaurosis, a congenital retinal blinding disease.^1,2^ The retina is an ideal candidate organ for gene-based therapies: it is readily accessible for surgical injection, can be imaged non-invasively in real-time, responds well to local treatments, and its blood-retinal-barrier limits unintentional spread of vectors to neighboring tissues and the general circulation.^3, 4^ Moreover, the retina is a post-mitotic tissue in which gene transfer induced by non-integrating vectors can achieve long-term production of therapeutic protein. Finally, the relatively immunoprivileged environment of the eye in general,^5^ and the retina in particular, may protect rAAV vectors from neutralization by pre-existing circulating anti-AAV antibodies. Clinical trials increasingly report adverse events in cohorts of patients injected with the highest rAAV doses.^6^ Inflammation and increased intraocular pressure were described in the high-dose cohort in a recent clinical trial of diabetic macular edema (DME, NCT04418427) patients. With the development of more effective viral vectors for retinal gene delivery, therapeutic thresholds could be reached with lower vector doses, thereby reducing the likelihood of the aforementioned adverse events.

The most common strategy to optimize rAAV vector efficacy involves modification of the viral capsid via genetic engineering, using either rational design or directed evolution approaches. Rational design approaches seek to introduce targeted changes in the capsid based on prior knowledge of capsid structure and function.^7,8^ By contrast, directed evolution approaches consist of repeated cycles of random mutations or peptide insertions guided by desired biological traits rather than existing knowledge.^9,10^ Although these strategies are promising,^11^ manufacture of new modified rAAV vectors is challenging, as it requires optimization of each stage of the production and purification processes, as well as complex characterization of the newly generated particles. We have developed a chemical engineering approach to rAAV vector optimization based on the presence of specific basic amino-acids on the AAV capsid. The use of a specific ligand could improve the targeting and transduction of specific cells but also modify the physico-chemical properties of the AAV, potentially increasing its efficacy. We previously described how functionalization of amino groups of rAAV2 capsids with a sugar bearing an isothiocyanate coupling function increases hepatocyte transduction *in vitro* with respect to nonmodified vectors.^12^ A significant advantage of this strategy is that we use “natural recombinant AAV vectors” meaning that no genetic modification is required to produce the starting product. In the present study, in order to specifically target retinal cells, we synthesized a new ligand, an arylisothiocyanate mannose derivative (**Figure 1**). This sugar was chosen because it can modify the physico-chemical properties of molecules or biomacromolecules after chemical bioconjugation by increasing their hydrosolubility. ^13^ We decided to use the AAV2 serotype based on our experience of its limited retinal diffusion, with transduction limited to the region of the bleb after subretinal injection. ^14^ As a proof of concept mannose grafting on the capsid of AAV2 may increase the diffusion and also the number of transduced cells. We assessed the efficacy of these chemically modified vectors following subretinal injections in rats and nonhuman primates. This new approach to the development of rAAV vectors has important implications for the rational design of safer and more effective AAV vectors for ophthalmic and other applications.

**Figure 1.**
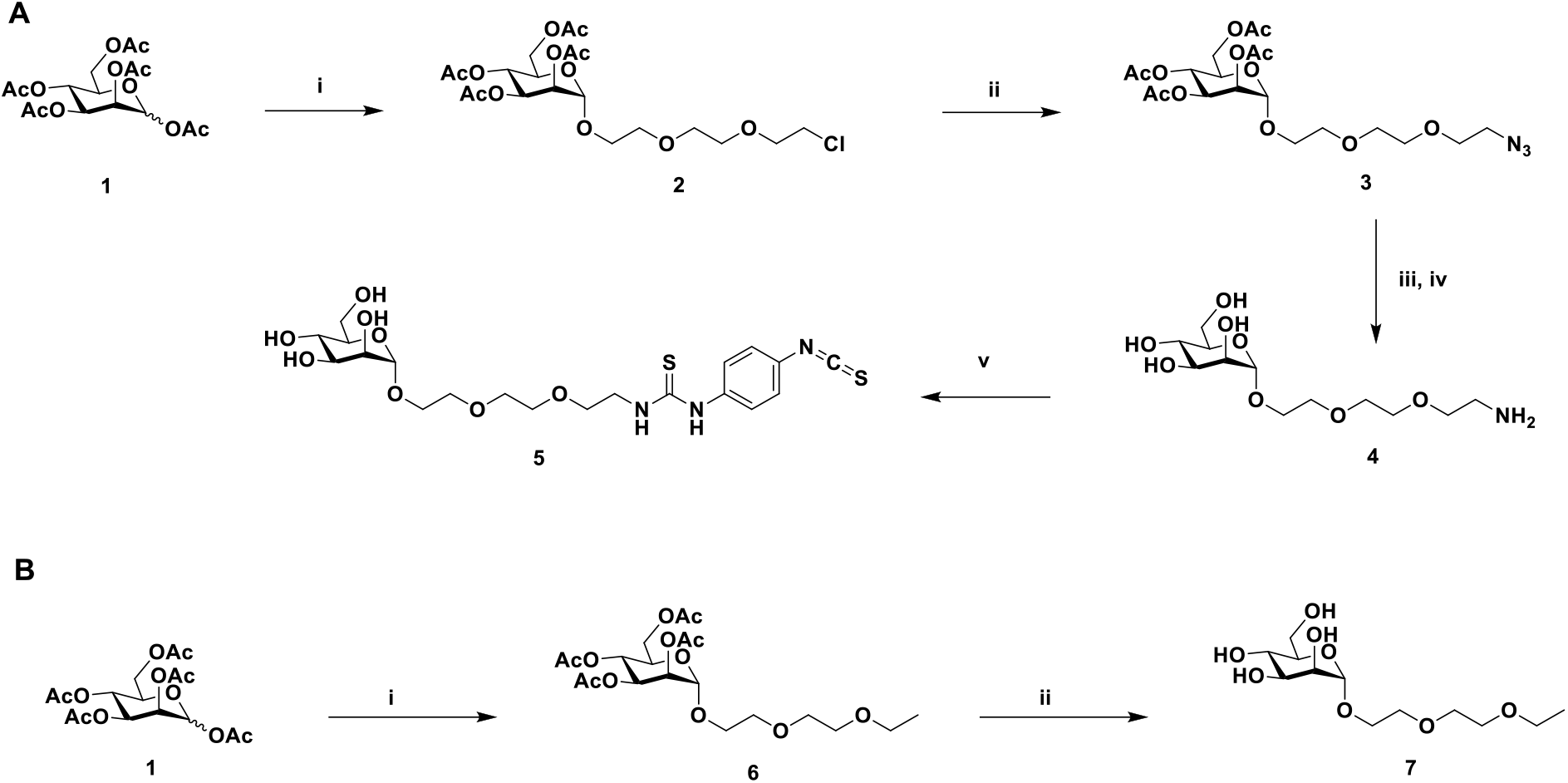
Synthesis of mannose derivatives with Ar-NCS coupling function and without the coupling function. (**A**) i: DCM, BF_3_.Et_2_O, 2-[2-(2-chloroethoxy)ethoxy]ethanol, molecular sieves (0°C 15 min then 8h RT, 73%), ii: DMF, NaN_3_ (16h, 70°C, 70%), iii: MeOH, APTS, H_2_, Pd/C (12h, RT, 99%), iv: MeOH/H_2_O, IRN78 (99%), v: DMF, *p*-phenylenediisothiocyanate (2h, RT, 44%). (**B**) i: DCM, BF_3_.Et_2_O, 2-(2-ethoxyethoxy)ethanol, molecular sieves (0°C 30 min then 12h RT, 72%), ii: MeOH/H_2_O, IRN78 (3h, RT, 99%).

## 2. Results and discussion

### 2.1 Synthesis of mannose ligands

We hypothesized that the development of rAAV particles carrying mannose carbohydrates would modifiy the physico-chemical properties of rAAV particles by increasing their solubilities and allows an increase of the transduced cells. Two sets of mannose ligands were designed and synthesized: one with a phenylisothiocyanate function capable of reacting with lysine residues bearing a primary amino group; and a second that lacked any reactive function and served as a control.

The chemical synthesis of the arylisothiocyanate (Ar-NCS) Compound **5** is shown in **Figure 1-A**. In the first step, the per-acetylated mannose **Compound 1** was glycosylated in the presence of BF_3_-Et_2_O and 2-[2-(2-chloroethoxy)ethoxy]-ethanol to obtain a significant yield of **Compound 2** in an exclusive α anomer configuration. The azide **Compound 3** was synthesized by nucleophilic substitution of the chloride of **Compound 2** in the presence of sodium azide and potassium iodide in dimethylformamide. This intermediate was reduced under hydrogen atmosphere with Pd/C in the presence of *p*-toluenesulfonic acid while deprotection of the hydroxyl groups was achieved using the basic resin IRN78 to obtain a quantitative yield of **Compound 4**. Finally, the Ar-NCS **Compound 5** was obtained by reacting **Compound 4** with an excess of *p-*phenylenediisothiocyanate in dimethylformamide with 44% yield. The Compound **7**, which lacked any coupling function, was synthesized in two steps from commercial D-mannose pentaacetate, with an overall yield of 71%. This control was synthesized to demonstrate that only covalent coupling on rAAV capsid surface occurs with ligand **5** and to control for any passive adsorption of the mannose-based compounds to the capsid surface during the chemical coupling. Glycosylation of C**ompound 1** in the presence of BF_3_-Et_2_O and 2-(2-ethoxyethoxy)-ethanol and cleavage of the acetyl groups resulted in the formation of the non-reactive Compound **7** (**Figure 1-B)**. All chemical compounds were characterized by NMR, HPLC, and mass spectrometry (**Figure S1-S12**).

### 2.2 Bioconjugation of mannose ligands to the AAV capsid

Bioconjugation of the mannose ligands **5** and **7** to the rAAV2 capsid surface was performed using an optimized version of a previously described protocol.^12^ Chemical coupling was achieved by nucleophilic addition of the amino group of the capsid proteins to the reactive isothiocyanate motif of the ligand compound **5**, yielding a thiourea linkage between the rAAV2 and the mannose ligand. Two different molar *ratios* of compound **5** (3^E^5 and 3^E^6) were used to evaluate (i) the possibility of modulating the number of mannose ligands on the rAAV capsid surface; and (ii) the impact of this modulation on the therapeutic index of these new vectors. Compound **7** was used only at the highest molar ratio (3^E^6) to demonstrate the absence of adsorption to the capsid surface (**Figure 2A**).

**Figure 2.**
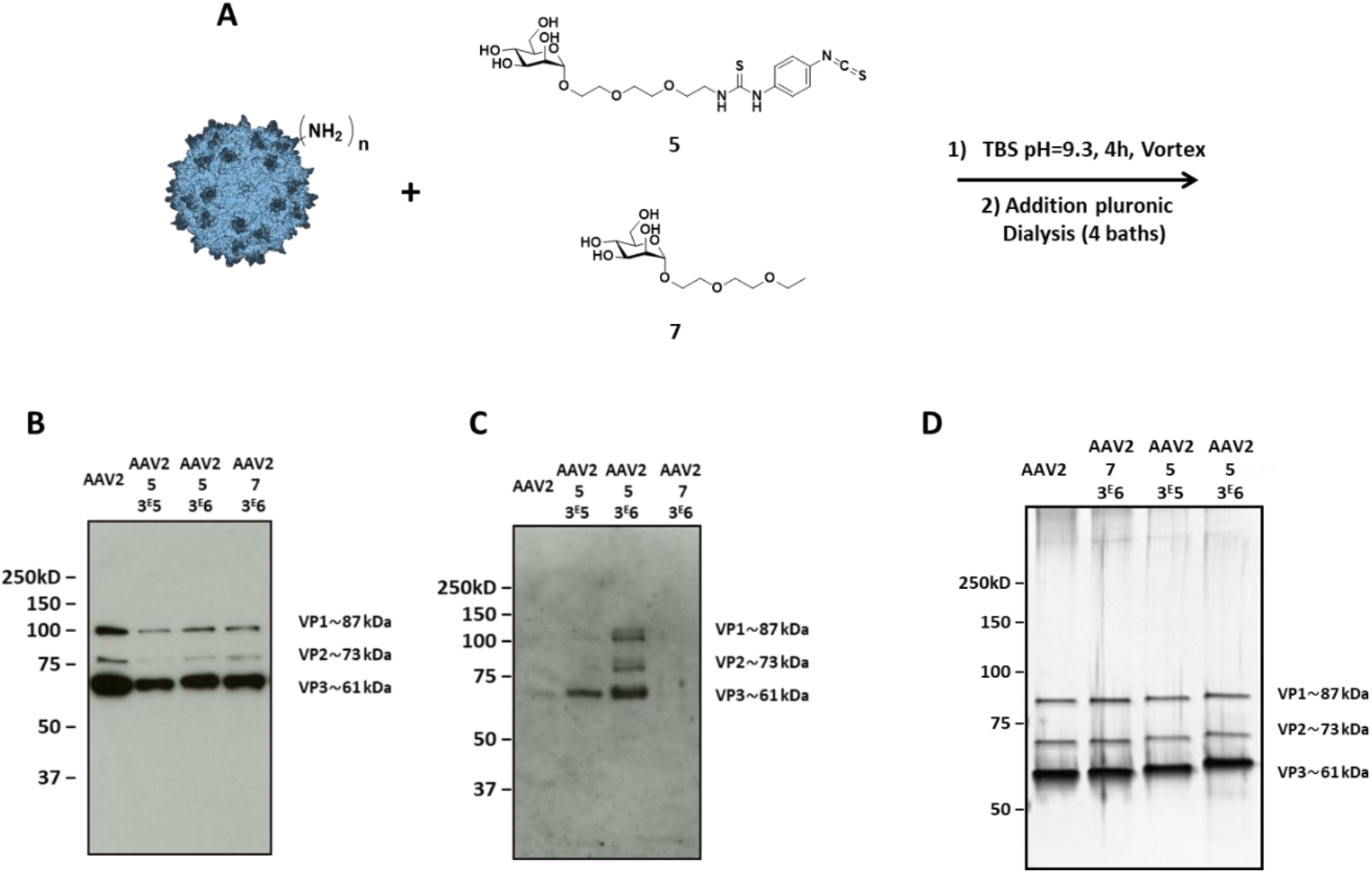
First covalent coupling of 5 onto the capsid of AAV2 *via* primary amino groups. (**A**) 1.4^E^12 vg of AAV2-GFP vectors were added to a solution of compound **5** (3^E^5 or 3^E^6 eq) in TBS buffer (pH 9.3) and incubated for 4 h at RT. The same experimental procedure was followed with compound **7** (3^E^6 eq) in TBS at pH 9.3 as a control. (**B, C**). 5^E^8 vg of the samples were analyzed by Western blot using a polyclonal antibody against the capsid to detect VP proteins (**B**) or using an FITC-concanavalin A lectin **(C)** to detect mannose sugar. Note that compound **7** is a negative control lacking the Ar-NCS reactive function. (**D**) 1^E^10 vg of each condition were analyzed by silver nitrate staining. VP1, VP2 and VP3 are the three proteins constituting the AAV capsid. Capsid protein molecular weight is indicated at the right of the images according to a protein ladder.

Western blot was first used to validate bioconjugation between rAAV2 and ligand **5**. Capsid-specific polyclonal antibody detection confirmed the integrity of the VP capsid subunits after the chemical reaction (**Figure 2B**) for all conditions tested. **Figure 2C** shows the efficient grafting of ligand **5** onto the rAAV2 capsid using concanavalin A lectin (which recognizes mannose sugar) demonstrating the covalent coupling of compound **5** onto the three AAV capsid proteins (VP1, VP2 and VP3).^15^ Conversely, rAAV2 capsid subunits incubated with ligand **7** could not be detected using concanavalin A lectin, confirming that the Ar-NCS function is essential for bioconjugation (**Figure 2C**). The amount of ligand **5** covalently attached to the three VPs clearly increased when the molar ratio was increased from 3^E^5 to 3^E^6 (**Fig. 2C)**, as reflected by the increase of the intensity of the bands. The purity and integrity of these mannosylated VPs was verified by silver staining, which showed that (i) VP1, VP2, and VP3 remained intact after the chemical coupling; and (ii) the molecular weight of each VP increased when the molar ratio was increased from 3^E^5 to 3^E^6 (**Figure 2D**).

Together these findings clearly demonstrate efficient bioconjugation of the mannose ligand **5** to the rAAV2 capsid via a thiourea covalent bond.

Aggregation of rAAVs is a well-described phenomenon.^16^ We therefore evaluated this parameter after bioconjugation. The size of rAAV2 and mannose-coupled rAAV2 particles was evaluated by dynamic light scattering (DLS) (**Table 1**). As shown in **Table 1**, the size of all particles ranged from 25–29 nm, indicating that the coupling of mannose ligands to the rAAV2 capsid did not promote aggregation.

**Table 1.** Size and infectivity of mannose-coupled AAV2 vectors. The size in nm was obtained by dynamic light scattering for each sample. Vector genome titers (vg/mL) were measured as indicated in the Materials and Methods section. The ratio of viral genomes to GFP forming units (vg/GFU) obtained in HEK (first coupling) and HeLa cells (second coupling) was evaluated. The lower this ratio, the more infectious the particles in this cell line.

|  | Samples | Titer<br>(vg/mL)<br>before<br>chemical<br>process | Titer<br>(vg/mL)<br>after<br>chemical<br>process | Size<br>(nm) | GFU/mL | vg/GFU |
| --- | --- | --- | --- | --- | --- | --- |
| **First**<br>**coupling** | **Unmodified**<br>**AAV2** | 1.7<sup>E13</sup> | - | 26.2 | 6.3<sup>E10</sup> | 1.0<sup>E2</sup> |
|  | **AAV2 + 5**<br>**(3<sup>E5</sup>)** | 1.4<sup>E12</sup> | 1.2<sup>E12</sup> | 25.6 | 5.6<sup>E9</sup> | 2.2<sup>E2</sup> |
|  | **AAV2 + 5**<br>**(3<sup>E6</sup>)** | 1.4<sup>E12</sup> | 1.3<sup>E12</sup> | 28.0 | 8.2<sup>E8</sup> | 1.6<sup>E3</sup> |
|  | **AAV2 + 7**<br>**(3<sup>E6</sup>)** | 1.4<sup>E12</sup> | 1.6<sup>E12</sup> | 26.0 | 6.6<sup>E9</sup> | 2.3<sup>E2</sup> |
| **Second**<br>**coupling** | **Unmodified**<br>**AAV2** | 1.0<sup>E13</sup> | - | 25.0 | 3.2<sup>E10</sup> | 3.2<sup>E2</sup> |
|  | **Dialyzed**<br>**AAV2** | 8<sup>E11</sup> | 7.7<sup>E11</sup> | 24.2 | 2.6<sup>E9</sup> | 3.0<sup>E2</sup> |
|  | **AAV2 + 5**<br>**(3<sup>E6</sup>)** | 8<sup>E11</sup> | 5.0<sup>E11</sup> | 29.0 | 1.1<sup>E7</sup> | 4.9<sup>E4</sup> |
|  | **AAV2 + 7**<br>**(3<sup>E6</sup>)** | 8<sup>E11</sup> | 5.5<sup>E11</sup> | 25.0 | 1.6<sup>E9</sup> | 3.1<sup>E2</sup> |

To demonstrate the reproducibility of the bioconjugation process, AAV coupling was repeated using new batches of rAAV2 and ligand **5** (3^E^6 eq). As shown in **Figure S13** and **Table 1**, all findings (qPCR, Western blot, silver staining, and DLS) were comparable with those obtained for the first batch, confirming the reproducibility of this technology.

As already described for rAAV2-GalNAc vectors, chemical modification of the capsid surface can impact on the infectivity of the modified particles.^12^ Indeed, chemical modification of lysine amino-acids on rAAV2 capsid motifs may limit or even abrogate targeting of the recombinant virus to heparan sulfate, one of its common receptors. Consequently, the tropism of the chemically-modified rAAV2 vectors could be altered, allowing transduction of cells that are usually non-permissive to rAAV2, while reducing the transduction of cells that express heparan sulfate receptors on their surface. This hypothesis is supported by the relative transduction efficiency observed for all mannose-coupled rAAV2 vectors carrying a GFP reporter gene when evaluated in HEK293 and HeLa cells. As shown in **Table 1**, vector genomes (vg) per GFP forming units (vg/GFU) values were in the same range for rAAV2, rAAV2 + **5** (3^E^5 eq) and rAAV2 + **7** (3^E^6 eq), and about one log higher for rAAV2 + **5** (3^E^6 eq). Comparable results were obtained for the two coupling experiments. The higher vg/GFU values obtained using rAAV2 coupled to ligand **5** (3^E^6 eq) may reflect reduced infectivity of these modified vectors: the higher ligand **5** payload may block the capsid motifs that are most efficiently recognized by the heparan sulfate receptors, thereby decreasing transduction efficiency in cells that bear these receptors. This decrease was not observed for the rAAV2 + **5** (3^E^5 eq) vector, perhaps because the lower coating density resulted in reduced blockade of capsid motifs.

### 2.3 *In vivo* evaluation in rats

To evaluate the *in vivo* efficacy of the mannose-coupled rAAV2-vectors for gene transfer in retina, two distinct batches of mannose-coupled rAAV2 vector were injected in two groups of rats (n=18 and 15, respectively) (**Table 2**). In both groups, control rats received subretinal injection of 2.5 μL vehicle (*i*.*e*. the vector formulation buffer). The experimental groups received 2.5 µL of either rAAV2 (at a dose of 1^E^12 vg/mL) or mannose-coupled rAAV2 (rAAV2 + **5** (3^E^5) or rAAV2 + **5** (3^E^6)) at doses of 1^E^12 or 4^E^11 vg/mL, respectively (total viral genomes injected per eye: 2.5^E^9 and 1^E^9 vg, respectively). For evaluation of the first batch of mannose-coupled rAAV2, rats received injections of unmodified and Mannose-coupled rAAV2 vector into the right and left eyes, respectively, and were monitored up to 4 weeks post-injection. For evaluation of the second batch of mannose-coupled rAAV2 vectors, rats received injections of vehicle, rAAV2, or mannose-coupled rAAV2 + **5** (3^E^6) in both eyes (n=5 animals per experimental condition). Eye fundoscopy was used for *in vivo* monitoring of GFP transgene expression post-injection at timepoints up to 6 weeks (**Table 2 and Figure 3**). Retinal structure was analyzed by spectral domain optical coherence tomography (SD-OCT). SD-OCT images of all eyes obtained before injection and immediately before sacrifice (i.e. 4- or 6-weeks post-injection) revealed no structural alterations in the retina, and preservation of the different retinal layers in all areas scanned (data not shown). This finding is in agreement with a previous report in which no significant toxicity was observed for rAAV vector doses in the same range as used in the present study.^17,18^ Fundoscopic examination of retinas from eyes injected with unmodified rAAV2 (n=28) or mannose-coupled rAAV2-vectors (rAAV2 + **5** [3^E^5], n=9; rAAV2 + **5** [3^E^6], n=19) revealed similar findings: GFP fluorescence was detected as early as 1 week post-injection, increased up to 6 weeks post-injection (the latest time point examined), and was limited to the retinal bleb induced by sub-retinal injection (**Figure 3**). Notably, subjective examination with the naked eye revealed a larger area of fluorescence and greater fluorescence intensity at all time points in retinas injected with rAAV2 + **5** (3^E^6) (n=19 eyes) versus those injected with identical dose of unmodified rAAV2 (n=28 eyes). This difference in fluorescence intensity was not observed in retinas injected with rAAV2 + **5** (3^E^5) (n=9). In line with *in vitro* data, this finding suggests that ligand density on the surface of rAAV vectors may influence vector transduction efficiency *in vivo*.

**Table 2.** Summary of *in vivo* studies. Abbreviations: vg/mL, vector genomes per mL; N/A, not applicable; IHC, immunohistochemistry; OF-OCT, ocular fundus; optical coherence tomography; VCN, vector genome copy numbers; mRNA, messenger RNA; WM, whole mount imaging. * 2.5 µL and 150 µL of each vector were injected into the retina of rats and NHPs, respectively.

|  | Animals | Injected eyes | Vectors | Vector concentration (vg/ml)* | *In vivo* and postmortem analysis |
| --- | --- | --- | --- | --- | --- |
| First coupling | Rat n=9 | Right | AAV2 | 1<sup>E12</sup> | OF-OCT, IHC |
|  |  | Left | AAV2 + 5 (3<sup>E5</sup>) | 1<sup>E12</sup> | OF-OCT, IHC |
|  | Rat n=9 | Right | AAV2 | 1<sup>E12</sup> | OF-OCT, IHC |
|  |  | Left | AAV2 + 5 (3<sup>E6</sup>) | 1<sup>E12</sup> | OF-OCT, IHC |
| Second coupling | Rat n=5 | Right | Vehicle | N/A | OF-OCT, VCN |
|  |  | Left | Vehicle | N/A | OF-OCT, mRNA levels |
|  | Rat n=5 | Right | AAV2 | 4<sup>E11</sup> | OF-OCT, VCN |
|  |  | Left | AAV2 | 4<sup>E11</sup> | OF-OCT, mRNA levels |
|  | Rat n=5 | Right | AAV2 + 5 (3<sup>E6</sup>) | 4<sup>E11</sup> | OF-OCT, VCN |
|  |  | Left | AAV2 + 5 (3<sup>E6</sup>) | 4<sup>E11</sup> | OF-OCT, mRNA levels |
|  | NHP 1 | Left | Vehicle | N/A | OF-OCT, WM |
|  | NHP 2 | Left | AAV2 | 4<sup>E11</sup> | OF-OCT, WM |
|  | NHP 3 | Left | AAV2 + 5 (3<sup>E6</sup>) | 4<sup>E11</sup> | OF-OCT, WM |

**Figure 3.**
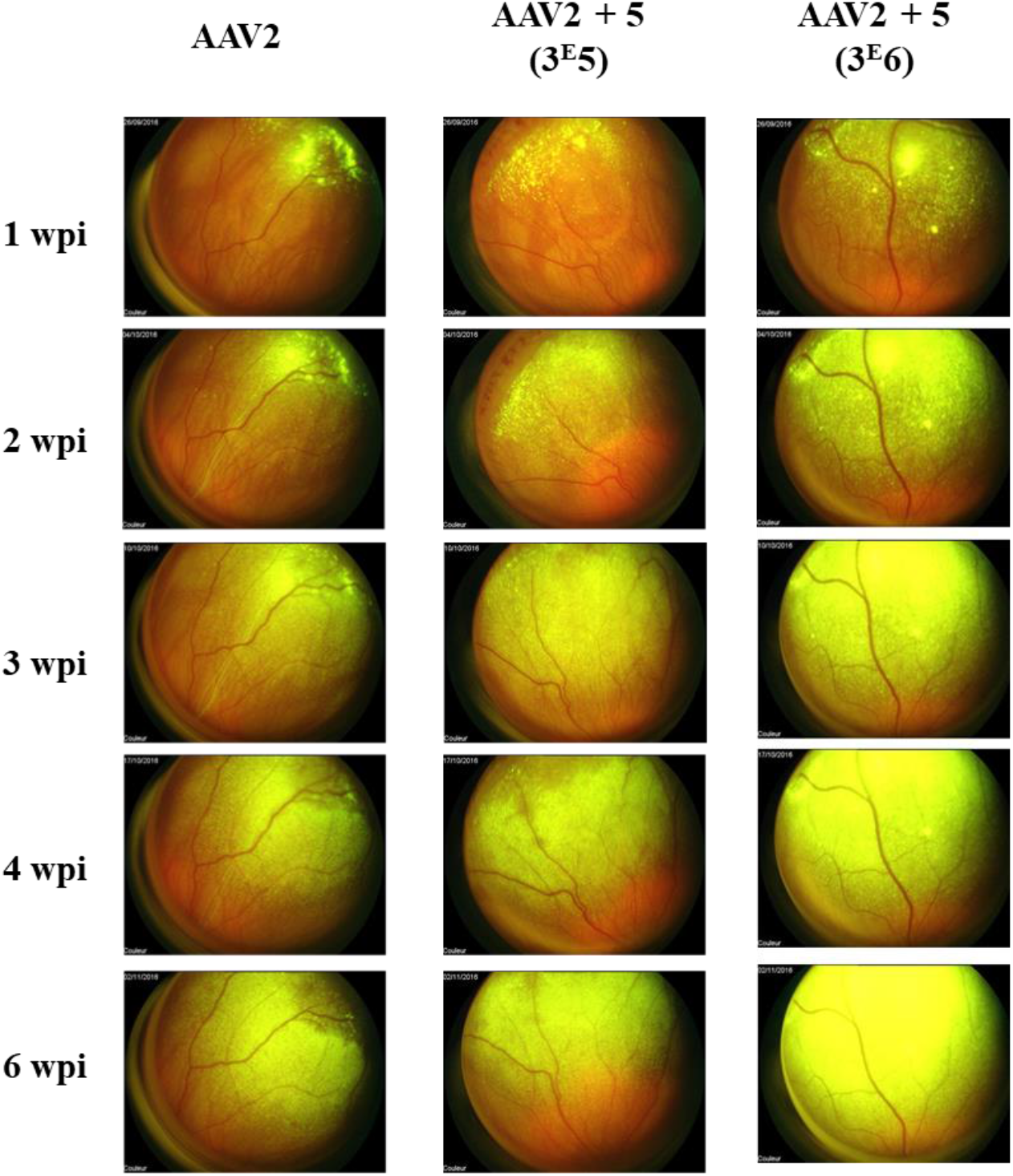
Ocular fundus in rats (first coupling). Kinetics of GFP expression after subretinal delivery of AAV2, AAV2 + **5** (3^E^5), or AAV2 + **5** (3^E^6), determined using *in vivo* imaging. Images show representative acquisitions obtained with each vector type. The green color becomes saturated on AAV2 + **5** (3e6) as the expression is very high, and the sensitivity of image acquisition on the retinophotograph is fixed for all retinal photographs from 1 wpi. Wpi, weeks post-injection.

To determine which cell types were transduced after subretinal injection of rAAV2 (n= 18 eyes) or mannose-coupled rAAV2 vector (rAAV2 + **5** [3^E^5], n=9 eyes; AAV2 + **5** [3^E^6], n=9 eyes), vertical sections were labeled with a fluorescent GFP antibody and overlayed with Mayer’s hemalum staining (**Figure 4**). Vertical sections were selected at random to determine the relative tropism of modified versus unmodified vectors, although it should be noted that this approach does not provide a quantitative measure of transduction (**Figure 4**).

**Figure 4.**
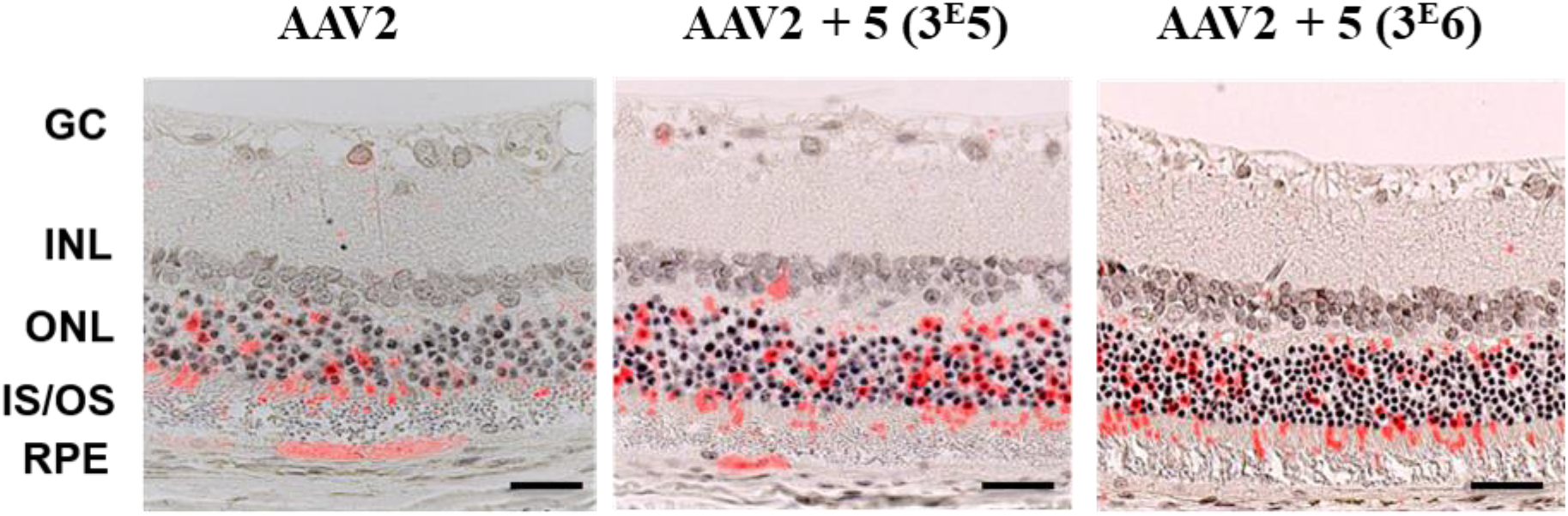
Vector distribution in the retina of rats. 4 weeks after subretinal injection with AAV2, AAV2+**5** (3E5), or AAV2+ **5** (3E6), from first coupling. Representative images of paraffin-embedded retina sections stained with anti-GFP antibody and anti-mouse Alexa 555-conjugated antibody (red labelling), and counterstained with hematoxylin. Scale bars, 30 *μ*m. GC, ganglion cells; INL, inner nuclear layer; ONL, outer nuclear layer; IS/OS, inner and outer segments; RPE, retinal pigment epithelium.

We observed that the same cell types were transduced by both modified and unmodified rAAV2 vectors: in both groups GFP was detected in the RPE and outer nuclear layer (ONL). Weaker transduction of the inner retinal layer and ganglion cells was observed. The cell tropism observed in our study is comparable to that previously described in rodent models injected subretinally with rAAV2 vectors.^19,20^

These findings indicate that chemical modification of rAAV2 with ligand **5** (3^E^6) did not modify rAAV2 tropism in the retina, but did result in higher transduction levels, which could have a beneficial impact on vector efficacy, thereby allowing for a reduction in vector dose. To investigate the basis of this apparent difference in transduction efficacy, we performed molecular biology analyses of retinas injected with unmodified and chemically modified rAAV2. We quantified vector copy number by qPCR (**Figure S15A-B**) and GFP mRNA transcripts by RT-qPCR (**Figure S16A-B**) 4 weeks post-injection in neuroretinas and the RPE from rats injected with vehicle (n=3), unmodified rAAV2 (n=5), or modified rAAV2 + **5** (3^E^6) (n=5) vector. In the neuroretina, vector genome copy numbers were significantly lower in rats injected with rAAV2 + **5** (3^E^6) than those injected with unmodified rAAV2 (mean 0.02 vg/dg and 0.10 vg/dg, respectively; p=0.0079). In the RPE, vector genome copy numbers were comparable regardless of the injection administered. However, in both the neuroretina and the RPE, GFP mRNA transcript levels were significantly (approximately 10-fold) higher in samples from rats injected with rAAV2 + **5** (3^E^6) than in those injected with unmodified rAAV2 (mean RQ=0.268 and 0.039, respectively in the neuroretina, p=0.0079; and mean RQ=0.244 and 0.017, respectively in the RPE, p = 0.0397). Next, in order to evaluate the “activity” of one vector genome copy, we calculated the ratio of GFP mRNA levels (relative quantity, RQ) to vector copy number (vector genomes per diploid genome, vg/dg) in the neuroretina (**Figure 5A**) and RPE (**Figure 5B**). In the neuroretina we observed significant differences between rats injected with unmodified rAAV2 vector and those injected with the modified rAAV2 + **5** (3^E^6) vector: a single copy of the rAAV2 + **5** (3^E^6) vector expressed up to 35 times more GFP mRNA than a single copy of the rAAV2 vector (**Figure 5A** – mean RQ/vg for rAAV2= 0.4, and mean RQ/vg for rAAV2 + **5** (3^E^6) = 13.6; p=0.0079). Note that this graph shows the ratio of transcript to vector genome. Hence, in a given retinal cell just one vector genome from the mannose-coupled vector can yield on average 17 transgene transcripts while it requires two vector genomes from the unmodified vector to have the probability of producing just one transcript. Similarly, in RPE samples GFP mRNA levels corresponding to a single copy of rAAV2 + **5** (3^E^6) vector were higher than those of the rAAV2 vector (mean RQ/vg for rAAV2=0.7, and mean RQ/vg for rAAV2 + **5** (3^E^6) = 4.8), although this difference was not significant, likely due to the high variability of RQ/vg ratios in rats injected with the mannose-coupled rAAV2 vector (**Figure 5B**).

**Figure 5.**
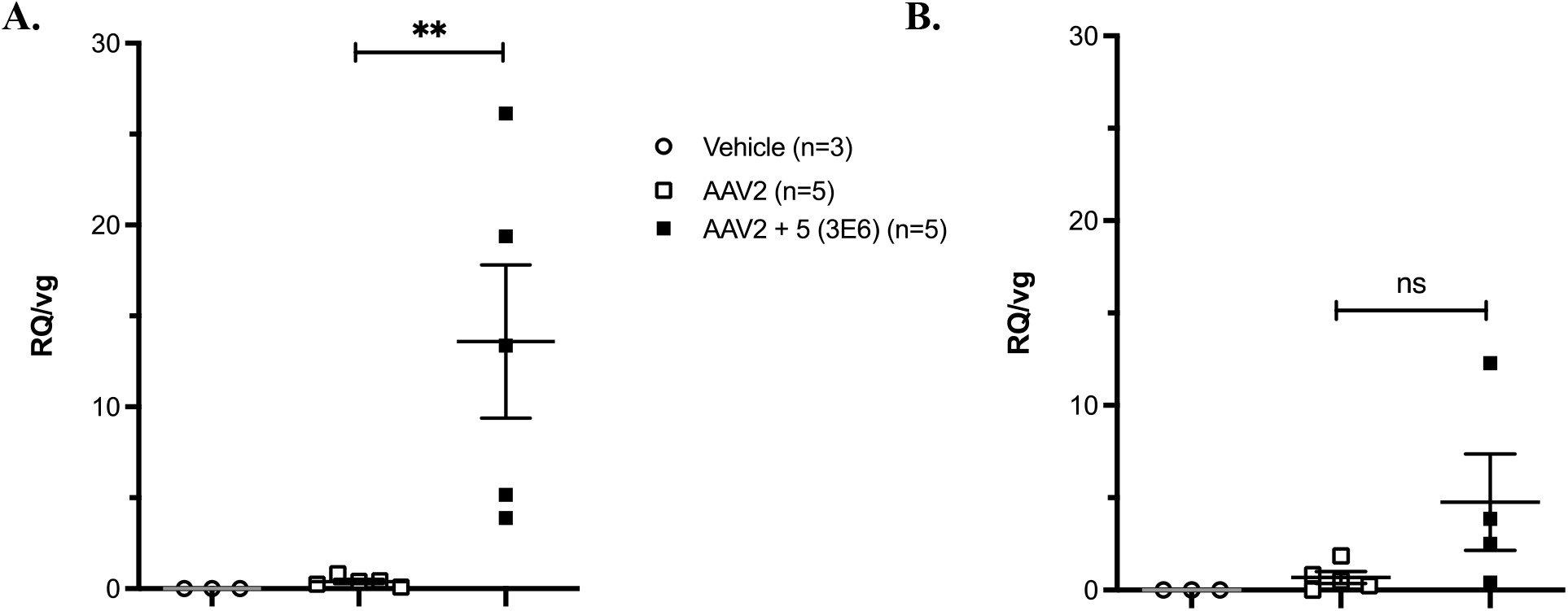
Individual ratio of GFP mRNA levels (Relative Quantity RQ) to vector copy number expressed as viral genome/diploid genome number (vg/dg), measured 4 weeks post-injection in the left and right neuroretinas (A) and in the RPE (B) of rats injected with vehicle, AAV2, or AAV2 + **5** (3^E^6) (second coupling). Results are expressed as mean ± SEM. Mann Whitney test was used for statistical analyses: ** p<0.01.

Taken together, these data showed that, compared with unmodified rAAV2, mannose-coupled rAAV2 vector exhibits similar cellular tropism but far superior transduction efficiency in the rat neuroretina.

### 2.4 *In vivo* evaluation in nonhuman primates (NHP)

Compared to rodents, the NHP retina has a thicker vitreous, a thicker inner limiting membrane, and a specialized fovea area with high visual acuity.^21^ We tested the efficacy of modified mannose-coupled rAAV2 vectors for retinal gene transfer in NHPs. Three (n=3) NHPs received subretinal injections of vehicle, rAAV2, or chemically modified mannose-coupled rAAV2 (rAAV2 + **5** [3^E^6]) in the left eye, between the RPE and photoreceptor layer (**Table 2**). Subretinal injections were performed using a transvitreal approach, following vitrectomy with air exchange, following the same surgical technique used for the administration of Luxturna®^22^ in patients with RPE65-mediated inherited retinal dystrophy. Surgery was well tolerated by all animals.

A vector concentration of 4^E^11 vg/mL was used, allowing for injection of a total viral load *per* eye of between 6^E^10 and 7^E^10, a dose conventionally used in preclinical and clinical settings.^22-24^ Animals were followed-up for 4 weeks after injection. Retinal structure was analyzed by SD-OCT before injection and immediately before euthanasia. No structural alterations or visible abnormalities were observed, and organization of the different retinal layers was preserved (data not shown). Expression of the GFP reporter was monitored by fundus fluorescence imaging before injection and at 1, 2, 3, and 4 weeks post-injection (**Figure 6**). As expected, no GFP fluorescence was observed in the eye injected with vehicle only. Fundoscopic examination of NHPs that received rAAV2 and rAAV2 + **5** (3^E^6) vector revealed GFP fluorescence as early as 1 week after vector delivery, after which a continuous increase in fluorescence was observed up to at least 4 weeks post-injection. For the same vector dose, we observed greater fluorescence intensity at each timepoint in the retina injected with the mannose-coupled rAAV2 vector, supporting our observations in the rat model. To evaluate the cellular tropism of rAAV2 and chemically modified rAAV2, retinas were flat-mounted and stained to label both cell nuclei (Draq5, in purple) and cone photoreceptors (LM and S opsin, in red) (**Figure 7**). The intensity of staining in all retinas treated with chemically modified rAAV2 was far greater than for the non-modified rAAV2 such that the direct comparison of the two groups cannot be made without saturation of the signals from the AAV2 + **5** (3E6) injected retinas. Nonetheless it was evident that there were no differences in the cell types transduced by the mannose-coupled rAAV2 vectors compared to unmodified rAAV2 vectors. Transduced cells included retinal ganglion cells, cells of the outer plexiform layer (OPL) and the outer nuclear layer (ONL), and photoreceptor inner segments. Almost no transduced cells were detected in the inner nuclear layer (INL) or in photoreceptor outer segments. This tropism was consistently observed across the entire transduced area of retina. In agreement with eye fundoscopy images showing greater fluorescence intensity in the NHP fundus, immunohistochemistry revealed higher GFP expression in retinal flat-mounts from NHPs injected with chemically modified mannose-coupled rAAV2 (**Figure 7B**). However, the proportion of the transduced area was similar in retinas injected with mannose-coupled rAAV2 vector (57%) and unmodified rAAV2 vector (60%) (**Figure 8**).

**Figure 6.**
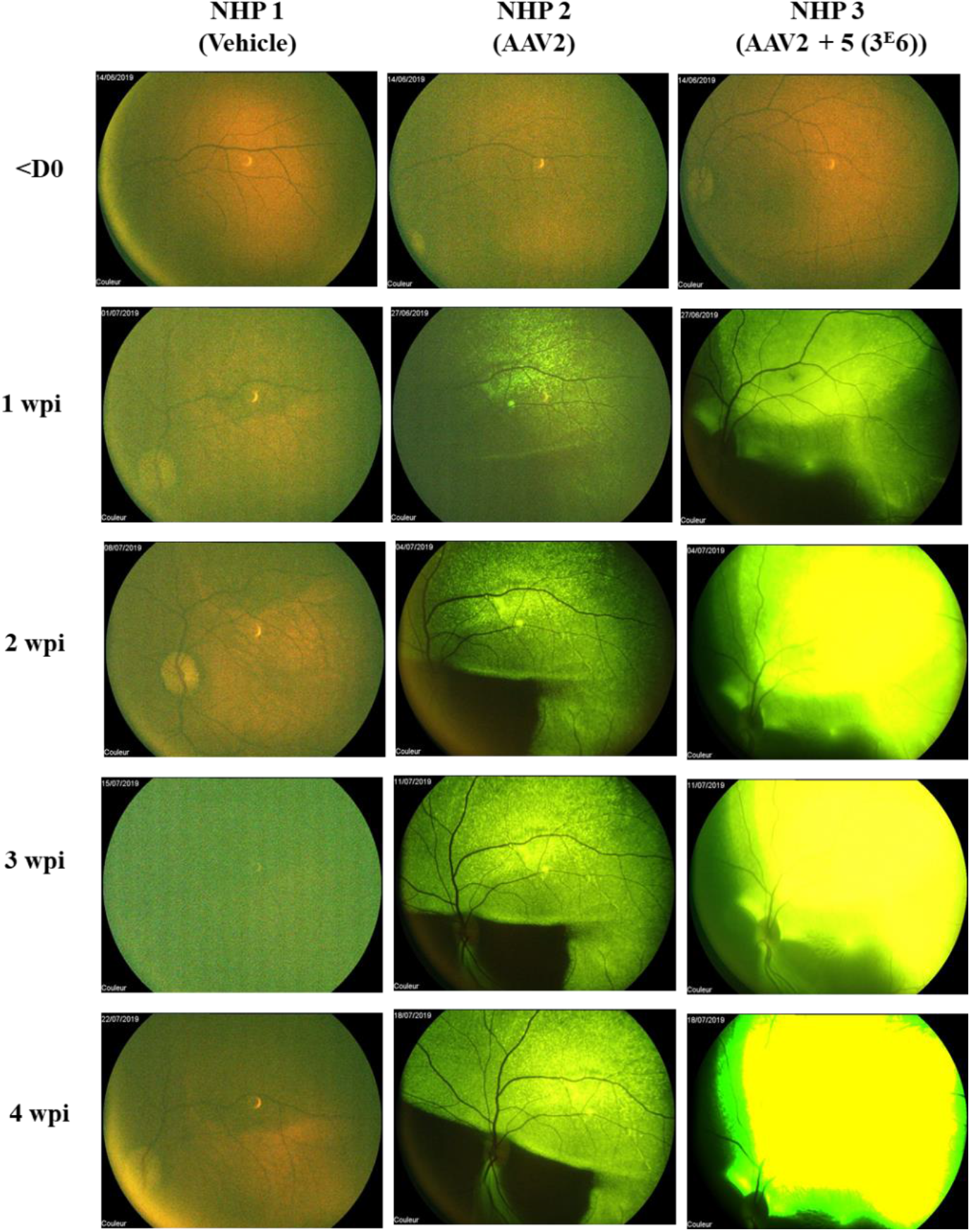
Ocular fundus in nonhuman primates. Kinetics of GFP expression after subretinal delivery of vehicle, AAV2, or AAV2+5 (3^E^6), using *in vivo* fundus imaging. The green color becomes quickly saturated for AAV2 + **5** (3e6), as the expression of GFP is very high, and the sensitivity of image acquisition on the retinophotograh was fixed for all retinal photographs from 1wpi. Note that retinal structure was preserved as normal on retinophotographs with lower sensitivity and as seen by OCT (data not shown). wpi, weeks post-injection.

**Figure 7.**
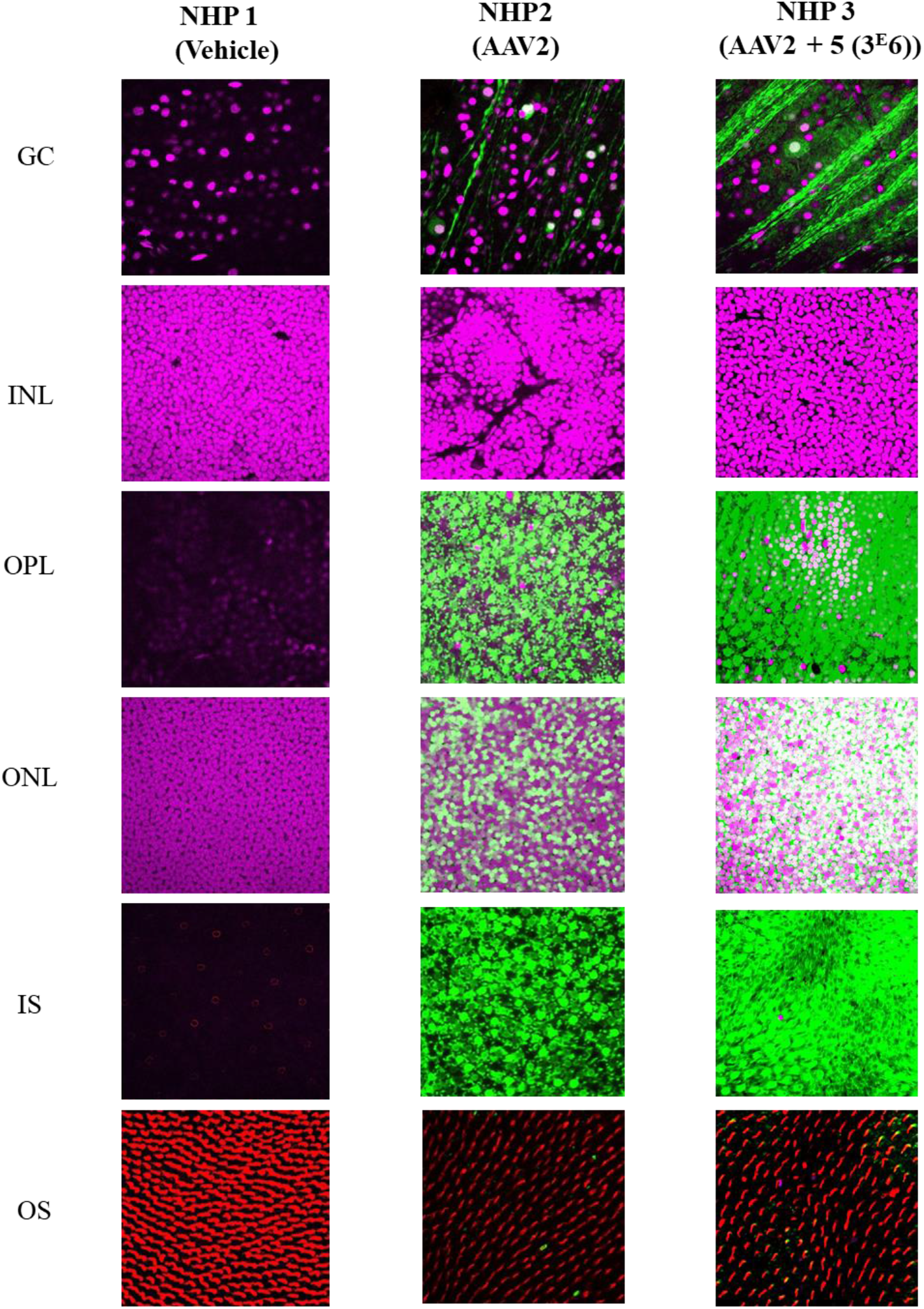
Retinal tropism of AAV2+5 (3^E^6) and AAV2 vectors in flat-mount retinas from nonhuman primates. Images are taken from confocal slices of each retinal layer and show GFP expression (green), nuclear labeling (purple), and immunostaining with LM opsin to identify outer segments (red). Original magnification ×60. Sensitivity of image acquisition was set on the retina with injection of AAV2, at the minimum to allow the visualization of the whole transduced retina. Images shown on this figure correspond to the temporal retina of both NHP2 and NHP3, where the GFP expression was maximal.

**Figure 8.**
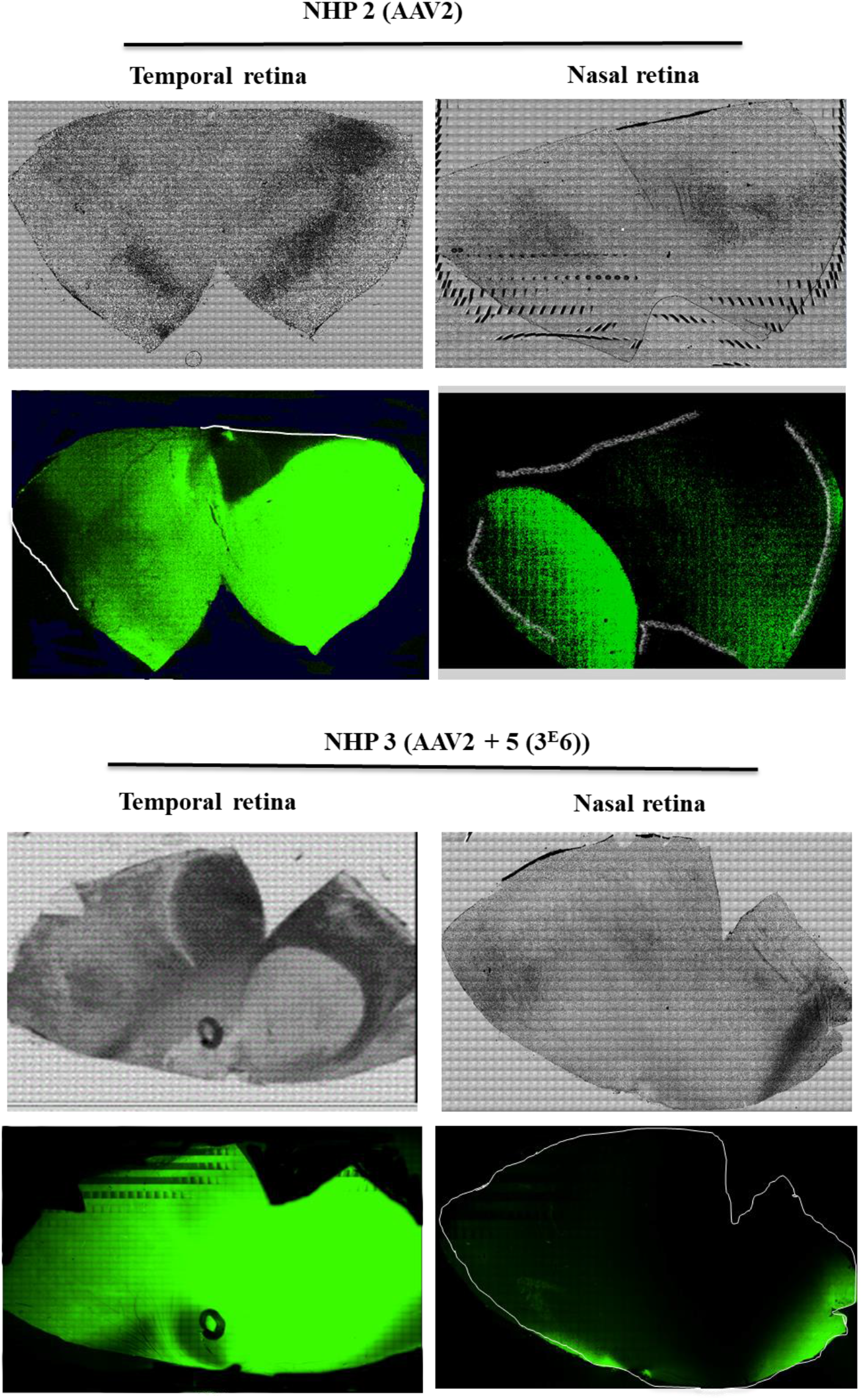
GFP expression in retinal whole mounts (nasal and temporal parts) from nonhuman primates following subretinal injection of AAV2 or AAV2+5 (3^E^6). Confocal images of whole-mounted retina allow estimation of the transduction area of AAV2 and AAV2 + 5 (3^E^6). Sensitivity of images acquisition was set on the retina with injection of AAV2, at the lowest setting to allow for the visualization of the whole transduced retina.

Given the increased retinal transduction efficiency of chemically-modified rAAV2, we used qPCR to compare the biodistribution in distant tissues (liver, spleen, occipital cortex, and optic chiasm) of the mannose-coupled rAAV2 vector versus unmodified rAAV2, following subretinal delivery. Low numbers of vector genome copies (0.002 vg/dg; LLOQ of our qPCR assay, 0.001 vg/dg) were detected only in the occipital cortex of the animal injected with rAAV2 + **5** (3^E^6). No vector genome copies were detected in any of the other samples from NHPs injected with rAAV2 or rAAV2 + **5** (3^E^6) vector. These results confirm previous reports describing an absence of unmodified rAAV biodistribution in the liver after subretinal delivery ^25,26^. Importantly, these data indicate that no extraocular distribution of the chemically modified mannose-coupled rAAV2 occurs after subretinal delivery in nonhuman primates.

## 3. Conclusion

In summary, we have demonstrated a novel biosynthetic approach to generate a new generation of chemically coupled enhanced gene therapy rAAV vectors. The novel rAAV2 vectors described here demonstrate altered transduction capacity both *in vitro* and *in vivo* through modification of the density of mannose ligands on the vector capsid surface. We show that subretinal delivery of mannose-coupled rAAV2 vectors into rat and NHP eyes does not change retinal cell tropism compared with unmodified rAAV2. Importantly, we observed significantly higher transgene expression in rat and NHP retinas injected with chemically modified mannose-coupled rAAV2 vector.

Taken together, our findings have important implications for ocular gene transfer studies and potential future clinical applications. We chose the AAV2 capsid to demonstrate proof-of principle in the serotype that is most well characterized across a range of tissues. However, other capsid types, both natural and engineered, could in principle be similarly chemically modified to increase transgene payload with the goal of reaching a therapeutic threshold with less virus. Reducing rAAV doses in the retina not only provides economic advantages by decreasing manufacturing costs, but could also reduce the risk of dose-related rAAV toxicity in treated patients, a phenomenon increasingly reported in clinical trials.

Moreover, altering the density of mannose ligands on the rAAV vector surface could have an impact on rAAV immunogenicity itself, with the induction of fewer anti-capsid neutralizing antibodies, as previously demonstrated in the liver for GalNac ligands on the surface of AAV vectors.^12^

In conclusion, our results demonstrate that modified mannose-coupled AAV2 vectors represent a new generation of biotherapeutic products for more efficient gene transfer in the retina application. This chemical modification of rAAVs could be applied to other serotypes relevant for ocular diseases to further improve gene therapy for retinal diseases.

## 4. Materials and methods

### 4.1 Materials

All chemical reagents were purchased from Acros Organics or Sigma Aldrich and were used without further purification. Rabbit polyclonal anti-AAV capsid proteins antibody (cat N° 61084) was obtained from PROGEN Biotechnik. Anti-fluorescein-AP Fab fragment antibody (cat N° 11426338910) for the detection of fluorescein-labeled lectin was obtained from Sigma-Aldrich. FITC-Concanavalin A lectin was purchased from Vector Laboratories. Reactions requiring anhydrous conditions were performed under nitrogen atmosphere. All chemically synthesized compounds were characterized by ^1^H (400º.133 or 300.135 MHz), ^13^C (125.773 or 75.480 MHz) NMR spectroscopy (Bruker Avance 300 Ultra Shield or Bruker Avance III 400 spectrometer). Chemical shifts are reported in parts per million (ppm); coupling constants are reported in units of Hertz [Hz]. The following abbreviations were used: s = singlet, d = doublet, t = triplet, q = quartet, quin = quintet, br = broad singlet. When needed, ^13^C heteronuclear HMQC and HMBC were used to unambiguously establish structures. High-resolution mass spectra (HRMS) were recorded with a Thermo Fisher hybrid LTQ-orbitrap spectrometer (ESI^+^) and a Bruker Autoflex III SmartBeam spectrometer (MALDI). HPLC analysis were performed on an HPLC autopurification system (WATERS) equipped with BGM 2545 binary pump, a 2767 Sample Manager, and an UV-visible diode array detector and evaporative light scattering detector in series (WATERS 996 PDA and WATERS 2424 ESLD). Chromatographic separation was performed on an Atlantis T3 (WATERS; 4.6 × 150 mm, 5µ). The mobile phase consisted of water (solvent A) and acetonitrile (solvent B). Gradient mix: 0.0 min [95% A; 5% B], 20.0 min [77% A; 23% B]. Flowrate: 1 mL/min.

All chemically synthesized products were purified by flash chromatography (GRACE REVELERIS Flash Chromatography System) equipped with UV and DLS detectors.

### 4.2 Synthesis

#### Compound 2

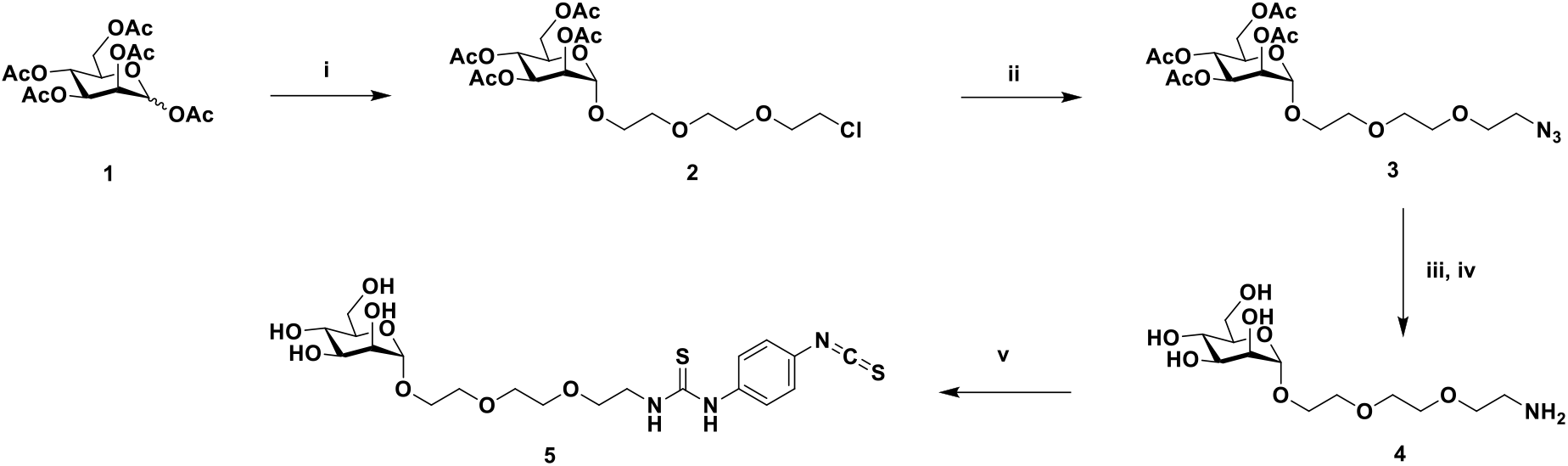

To a solution of peracetylated mannose **1** (2 g, 5.128 mmol, 1 eq) in dry DCM (51 mL) containing oven-dried powdered 4 Å molecular sieves, 2-[2-(2-chloroethoxy)ethoxy]ethanol (2.99 g, 7.692 mmol, 1.5 eq) was added and stirred at 0°C under N_2_ atmosphere for 15 min. F_3_B·OEt_2_ (1.90 mL, 15.384 mmol, 3 eq) was then added dropwise and continually stirred for 8 h at room temperature (RT). The mixture was diluted with DCM and washed with an aqueous saturated solution of NaHCO_3_. The crude of the mannosyl glycoside **2** was used in the next step without further purification (2.78 g).

#### Compound 3

To a solution of **the crude Compound 2** in dry DMF (30 mL), NaN_3_ (1.33 g, 20.512 mmol, 4 eq) was added and the mixture was stirred at 70°C under N_2_ atmosphere for 16 h. DMF was evaporated under vacuum conditions, and the residue was dissolved in AcOEt and washed with water and brine. The crude product was purified by column chromatography (SiO_2_, Cy/AcOEt, 70/30 as eluent) to produce the azide derivative **Compound 3** (1.81 g, 3.581 mmol, 70% in two steps), a colourless oil. ^1^H NMR (CDCl_3_): 1.98 (s, 3H, AcO), 2.03 (s, 3H, AcO), 2.10 (s, 3H, AcO), 2.15 (s, 3H, AcO), 3.39 (t, 2H, *J*_2,1_ = 5.1 Hz, **CH**_**2**_N_3_), 3.62-3.81 (m, 10H, 5 × **CH**_**2**_O), 4.03-4.12 (m, 2H, H-5, H-6a), 4.29 (dd, 1H, H-6b, *J*_6b,6a_ = 12.4 Hz, *J*_6b,5_ = 5.2 Hz), 4.87 (d, 1H, H-1, *J*_1,2_ = 1.7 Hz), 5.26 (dd, 1H, H-2, *J*_2,3_ = 3.2 Hz, *J*_2,1_ = 1.7 Hz), 5.28 (t, 1H, H-4, *J*_4,3_ = *J*_4,5_ = 9.9 Hz), 5.35 (dd, 1H, H-3, *J*_3,4_ = 9.9 Hz, *J*_3,2_ = 3.3 Hz); ^13^C NMR (CDCl_3_): 20.61 (CH_3_), 20.63 (CH_3_), 20.7 (CH_3_), 20.8 (CH_3_), 50.7 (CH_2_), 60.4 (CH_2_), 66.2 (CH), 67.4 (CH_2_), 68.4 (CH), 69.1 (CH), 69.6 (CH), 70.0 (2 × CH_2_), 70.7 (CH_2_), 70.8 (CH_2_), 97.7 (CH), 169.7 (C), 169.8 (C), 169.9 (C), 170.6 (C); HRMS (ESI) for C_20_H_31_O_12_N_3_Na [M+Na]^+^, calcd 528.1805 found 528.1805 (**Figs. S1 and S2**).

#### Compound 4

To a solution of **Compound 3** (225 mg, 0.445 mmol) in MeOH (4.5 mL), PTSA (77 mg, 0.449 mmol) was added, followed by 10% Pd-C (10% w). The resulting suspension was stirred under H_2_ atmosphere (1 atm) for 12 h. The Pd/C was removed by filtration through Celite^®^ and the filtrate was evaporated under reduced pressure to produce the ammonium salt, as confirmed by ^1^H NMR.

The crude of the reaction was dissolved in a 1:1 mixture of H_2_O/MeOH (20 mL), to which Amberlite IRN78 basic resin was added. After 3 h of stirring at 20°C, the reaction mixture was filtrated and evaporated under reduced pressure to yield deprotected amine **Compound 4**, a colorless oil (137 mg, 0.440 mmol, 99%), which was used in the next step without further purification. ^1^H NMR (MeOD): 2.81 (m, 2H, **CH**_**2**_NH_2_), 2.15-3.9 (m, 16H, 5 × **CH**_**2**_O, H-2, H-3, H-4, H-5, H-6a, H-6b), 4.81 (d, 1H, H-1, *J*_1,2_ = 1.7Hz); ^13^C NMR (MeOD): 41.9 (CH_2_), 62.9 (CH_2_), 67.7 (CH), 68.6 (CH_2_), 71.3 (CH_2_), 71.4 (CH_2_), 71.6 (CH), 72.1 (CH), 72.5 (CH), 72.8 (CH_2_), 74.6 (2 × CH_2_), 101.7 (CH); HRMS (ESI) for C_12_H_26_NO_8_ [M+H]^+^, calcd 312.1658 found 312.1651 (**Figs. S3 and S4**).

#### Compound 5

To a solution of **Compound 4** (138 mg, 0.444 mmol) in dry DMF (4.0 mL), *p*-phenylene diisothiocyanate (426 mg, 0.219 mmol, dissolved in 4.0 mL of dry DMF) was added dropwise under N_2_ atmosphere using a syringe pump over the course of 1 h. After 2 h of stirring at 20°C, the solvent was evaporated under reduced pressure and the residue was purified by flash chromatography (SiO_2_, DCM/MeOH: 100/0 → 80/20) to yield the thioisocyanate **Compound 5** (98 mg, 0.195 mmol, 44%), a light-yellow solid. ^1^H NMR (DMSO-d6): 3.29-3.72 (m, 18H, 5 × **CH**_**2**_O, **CH**_**2**_NH, H-2, H-3, H-4, H-5, H-6a, H-6b), 4.45 (t, 1H, OH, *J*_OH,H_ = 5.9 Hz), 4.58 (d, 1H, OH, *J*_OH,H_ = 5.9 Hz), 4.64 (d, 1H, H-1, *J*_1,2_ = 1.4 Hz), 4.73 (d, 1H, OH, *J*_OH,H_ = 4.5 Hz), 4.76 (d, 1H, OH, *J*_OH,H_ = 4.7 Hz), 7.37 (d, 2H, *J*_H,H_ = 8.9 Hz), 7.59 (d, 2H, *J*_H,H_ = 8.9 Hz), 7.93 (bs, 1H, NH), 9.79 (bs, 1H, NH); ^13^C NMR (DMSO-d6): 43.6 (CH_2_), 61.3 (CH_2_), 65.7 (CH_2_), 67.0 (CH), 68.5 (CH_2_), 69.5 (CH_2_), 69.6 (CH_2_), 69.8 (CH_2_), 70.3 (CH), 70.9 (CH), 73.9 (CH), 100.0 (CH), 123.1 (2 × CH), 124.7 (C), 126.2 (2 xCH), 132.6 (C), 139.3 (C), 180.3 (C); HPLC: t_R_ = 8.17 min, mobile phase consisting of water (solvent A) and acetonitrile (solvent B). Gradient mix: 0.0 min [95% A; 5% B], 20.0 min [77% A; 23% B]. Flowrate: 1 mL/min.; HRMS (ESI) for C_20_H_29_N_3_O_8_S_2_Na [M+Na]^+^, calcd 526.1294 found 526.1292 (**Figs. S5, S6, and S7**).

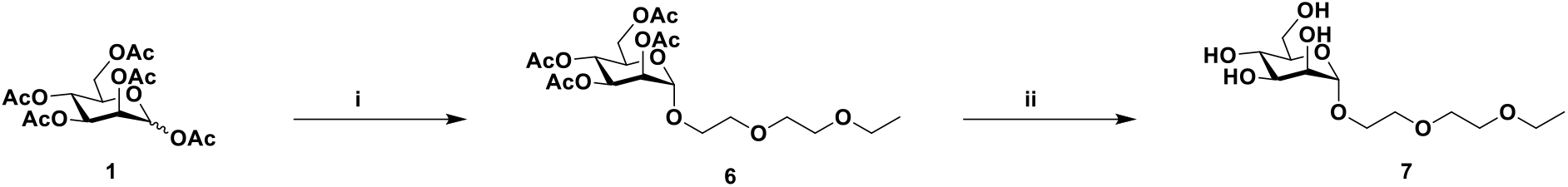

#### Compound 6

To a solution of the acetyl glycoside, **Compound 1** (500 mg, 1.282 mmol) and 2-(2-ethoxyethoxy)ethanol (258 mg, 1.923 mmol) in dry DCM (13 mL), 4 Å MS, was added. The resulting solution was stirred at 20°C under N_2_ atmosphere. After 30 min, F_3_B·OEt_2_ (791µL, 6.410 mmol) was added dropwise at 0°C under N_2_ atmosphere. The reaction mixture was stirred for 30 min at 0°C and warmed to 20°C. After 12 h of stirring, the MS was filtered and the solvent evaporated. The residue was redissolved in DCM, washed respectively with saturated aqueous NaHCO_3_, water, and brine, dried over MgSO_4_, filtered, and evaporated. The residue was purified by flash chromatography (SiO_2_, Cy/AcOEt 70/30) to yield the derivative **Compound 7** (430 mg, 0.926 mmol, 72%), a colorless oil. ^1^H NMR (CDCl_3_): 1.20 (t, 3H, **CH**_**3**_CH_2_O, *J* = 7.0 Hz), 1.97 (s, 3H, AcO), 2.02 (s, 3H, AcO), 2.09 (s, 3H, AcO), 2.14 (s, 3H, AcO), 3.52 (q, 2H, CH_3_**CH**_**2**_O, *J* = 7.0 Hz), 3.56-3.86 (m, 8H, 4 × **CH**_**2**_O), 4.08 (m, 2H, H-5, H-6a), 4.28 (dd, 1H, H-6b, *J*_6b,6a_ = 12.6 Hz, *J*_6b,5_ = 5.3 Hz), 4.89 (d, 1H, H-1, *J*_1,2_ = 1.5 Hz), 5.25 (dd, 1H, H-2, *J*_2,3_ = 3.3 Hz, *J*_2,1_ = 1.8 Hz), 5.27 (t, 1H, H-4, *J*_4,3_ = *J*_4,5_ = 9.7 Hz), 5.35 (dd, 1H, H-3, *J*_3,4_ = 10.0 Hz, *J*_3,2_ = 3.4 Hz); ^13^C NMR (CDCl_3_): 15.1 (CH_3_), 20.6 (CH_3_), 20.7 (CH_3_), 20.8 (CH_3_), 62.4 (CH_2_), 66.1 (CH), 66.6 (CH_2_), 67.4 (CH), 68.4 (CH), 69.1 (CH), 69.6 (CH_2_), 69.8 (CH_2_), 70.0 (CH_2_), 70.8 (CH_2_), 97.7 (CH), 169.7 (C), 169.8 (C), 170.0 (C), 170.1 (C); HRMS (ESI) for C_20_H_32_O_12_Na [M+Na]^+^, calcd 487.1791 found 487.1784 (**Figs. S8 and S9**).

#### Compound 7

The acetylated sugar **Compound 6** (310 mg, 0.667 mmol) was dissolved in a 1:1 mixture of H_2_O/MeOH (12 mL), to which Amberlite IRN78 basic resin was added. After 3 h of stirring at 20°C, the reaction mixture was filtrated and evaporated under reduced pressure to yield the final **Compound 7** (195 mg, 0.659 mmol, 99%), a white solid. ^1^H NMR (MeOD): 1.19 (t, 3H, **CH**_**3**_CH_2_O, *J* = 7.0 Hz), 3.54 (q, 2H, CH_3_**CH**_**2**_O, *J* = 7.0 Hz), 3.56-3.86 (m, 14H, 4 × **CH**_**2**_O, H-2, H-3, H-4, H-5, H-6a, H-6b), 4.79 (d, 1H, H-1, *J*_1,2_ = 1.7 Hz); ^13^C NMR (MeOD): 15.4 (CH_3_), 62.9 (CH_2_), 67.6 (CH), 67.8 (CH_2_), 68.6 (CH), 70.9 (CH), 71.4 (CH), 71.6 (CH_2_), 72.1 (CH_2_), 72.5 (CH_2_), 74.6 (CH_2_), 101.7 (CH); HPLC: t_R_ = 2.19 min, mobile phase consisting of water (solvent A) and acetonitrile (solvent B). Gradient mix: 0.0 min [95% A; 5% B], 20.0 min [77% A; 23% B]. Flowrate: 1 mL/min.; HRMS (ESI) for C_12_H_24_O_8_Na [M+Na]^+^, calcd 319.1369 found 319.1371 (**Figs. S10, S11, and S12**).

### 4.3 AAV2 production and purification

AAV2 vectors were produced from two plasmids: (i) pHelper, PDP2-KANA encoding AAV Rep2-Cap2 and adenovirus helper genes (E2A, VA RNA, and E4); and (ii) the pVector ss-CAG-eGFP containing the ITRs. All vectors were produced by transient transfection of HEK293 cells using the calcium phosphate-HeBS method. AAV2 transfected cells were harvested 48 h after transfection and treated with Triton-1% and benzonase (25 U/mL) for 1 h at 37°C. The resulting bulk was subjected to freeze-thaw cycles to release vector particles. The cellular debris were removed by centrifugation at 2500 rpm for 15 min. Cell lysates were precipitated with PEG overnight and clarified by centrifugation at 4000 rpm for 1 h. The precipitates were then incubated with benzonase for 30 min at 37 °C and collected after centrifugation at 10,000 g for 10 min at 4°C. Vectors were purified by double cesium chloride (CsCl) gradient ultracentrifugation. The viral suspension was then subjected to four successive rounds of dialysis with mild stirring in a Slide-a-Lyzer cassette (Pierce) against dPBS (containing Ca^++^ and Mg^++^).

### 4.4 Coupling and purification

AAV2-GFP (1^E^12 vg, 2.49 nmol) were added to a solution of TBS buffer (pH 9.3) containing **Compounds 5** or **7** at different molar ratios (3^E^5 or 3^E^6 equivalent), as stated in the respective results sections, and incubated for 4 h at RT. The solutions containing the vectors were then dialyzed against dPBS + 0.001% Pluronic to remove free molecules that had not bound to the AAV capsid.

### 4.5 Titration of AAV vector genomes

A total of 3 µL of AAV was treated with 20 units of DNase I (Roche #04716728001) at 37°C for 45 min to remove residual DNA in vector samples. After treatment with DNase I, 20 µL of proteinase K (20 mg/mL; MACHEREY-NAGEL # 740506) was added and the mixture incubated at 70°C for 20 min. An extraction column (NucleoSpin®RNA Virus) was then used to extract DNA from purified AAV vectors.

Quantitative real time PCR (qPCR) was performed with a StepOnePlus™ Real-Time PCR System Upgrade (Life Technologies). All PCRs were performed with a final volume of 20 µL, including primers and probes targeting the ITR2 sequence, PCR Master Mix (TaKaRa), and 5 µL of template DNA (plasmid standard or sample DNA). qPCR was carried out with an initial denaturation step at 95°C for 20 seconds, followed by 45 cycles of denaturation at 95°C for 1 second and annealing/extension at 56°C for 20 seconds. Plasmid standards were generated with seven serial dilutions (containing 10^8^ to 10^2^ plasmid copies), as described by D’Costa *et al*.^27^

### 4.6 Western blot and silver staining

All vectors were denatured at 100°C for 5 min using Laemmli sample buffer and separated by SDS-PAGE on 10% Tris-glycine polyacrylamide gels (Life Technologies). Precision Plus Protein All Blue Standards (BioRad) were used as a molecular-weight size marker. After electrophoresis, gels were either silver stained (PlusOne Silver Staining Kit, Protein; GE Healthcare) or transferred onto nitrocellulose membranes for Western blot. After transferring the proteins to nitrocellulose membrane using a transfer buffer (25 mM Tris/192 mM glycine/0.1 (w/v) SDS/20% MeOH) for 1 h at 150 mA in a Trans-Blot SD Semi-Dry Transfer Cell (Bio-Rad), the membrane was saturated for 2 h at RT with 5% semi-skimmed milk in PBS-Tween (0.1%) or with 1% gelatin, 0.1% Igepal in PBS-Tween (0.01%). After saturation, the membrane was probed with the corresponding antibody (anti-capsid polyclonal, B1 monoclonal, or anti-fluorescein-AP) and FITC-Concanavalin A lectin (mannose detection) overnight at 4°C. Three washes (15 min at RT) with PBS-Tween (0.1%) were performed between each stage to remove unbound reagents. Bands were visualized by chemiluminescence using alkaline phosphatase (AP) or horseradish peroxidase (HRP)-conjugated secondary antibodies and captured on X-ray film.

### 4.7 Dynamic light scattering

Dynamic light scattering (DLS) was performed using a Malvern Zetasizer Nano ZS. Calibration was controlled beforehand using a 30 and 300 nm solution of Nanosphere Size Standard. A volume of 50 µL of each vector was placed in a specific cuvette (DTS0118; Malvern) and analyzed by volume.

### 4.8 AAV infectious titer measurement

The infectivity of each sample was measured as follows. HEK293 or HeLa cells were seeded in 2 mL DMEM growth medium in 6-well culture plates at a density of 10^6^ cells/well. Cells were then incubated overnight at 37°C to reach 50% confluence. The viral stock was then diluted 10-fold by serial dilution. Next, 2 µL of each dilution was added to separate wells in the 6-well plates. Plates were then incubated at 37°C for 24 h. The infectivity of the AAV2-GFP control was measured immediately upon thawing of the sample. The same procedure was used for mannose particles. AAV-GFP-infected cells were detected by fluorescence microscopy.

The transducing unit (TU) titer was calculated using the following formula:

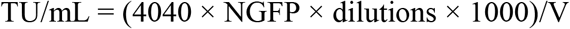

where NGFP is the mean number of GFP-positive cells per well and V is the volume (in µL) of vector used to infect cells.

### 4.9 Animal care and welfare

Experiments were performed on 33 rats (Sprague Dawley) and 3 male cynomolgus monkeys *(Macaca fascicularis*) provided by BioPrim (Baziege, France). Animals were euthanized 1 month after AAV injection. Research was conducted at the Boisbonne Centre (ONIRIS, Nantes-Atlantic College of Veterinary Medicine, Nantes) under authorization #H44273 from the Departmental Direction of Veterinary Services (Loire-Atlantique, France). All animals were handled in accordance with the Guide for the Care and Use of Laboratory Animals. Experiments involving animals were conducted in accordance with the guidelines of the Animal Experimentation Ethics Committee of Pays de Loire (France) and the Ministry of Higher Education and Research. Animals were sacrificed by intravenous injection of pentobarbital sodium (Dolethals; Vetoquinol) in accordance with approved protocols.

### 4.10 Subretinal injections

#### Injection in rats

Rats were anesthetized by inhalation of isoflurane gas followed by intramuscular injection of ketamine/xylazine (Imalgène1000/Rompun 2%). Pupils were dilated with 0.5% tropicamide and 10% neosynephrine eye drops. Local anesthesia was achieved using 0.4% oxybuprocaïne eye drops. The ocular surface was disinfected with 5% vetedine and rinsed 3 times with Ocryl®. Under an operating microscope a trans-scleral/transchoroidal tunnel incision was made using surgical nylon suture (Ethilon 10/0) and the AAVs delivered to the subretinal space by injection using a 33G needle and a 5-mL Hamilton syringe. Slow injection was performed at a rate of 2 μL/min while monitoring using a binocular microscope. A volume of 2.5 μL was injected with 1/1000 of fluorescein dye to monitor bleb size. Injection success was confirmed by fundus autofluorescence imaging immediately after injection by SD-OCT (Spectralis® HRA+OCT imaging system; Heidelberg Engineering Inc.).

#### Injection in nonhuman primates

Nonhuman primates were anesthetized by isoflurane gas inhalation. Pupils were dilated with atropine 0.3%, tropicamide 0.5%, and neosynephrine 10% eye drops. Local anesthesia was achieved with oxybuprocaine 0.4% eye drops. The ocular surface was disinfected with vetedine 5% and rinsed three times with Ocryl®. After lateral canthotomy, a 25-gauge 3-port vitrectomy was performed with posterior vitreous detachment (Vitreotom Accurus; Alcon France). Retinal bleb formation was induced by subretinal injection of viral vector (150 µL) along the temporal superior vascular arcade with a 41-gauge subretinal injection cannula. At the end of the procedure, fluid-air exchange of the vitreous cavity was performed. Sclerectomies and the lateral canthotomy were sutured with Vicryl® 7/0 (absorbable). Subconjunctival injection of 0.5 mL methylprednisolone (40 mg/mL) was performed.

### 4.11 Retinal imaging

For evaluation of retinal structure, SD-OCT was performed before AAV injection and after euthanasia using the Spectralis^®^ HRA+OCT imaging system (Heidelberg Engineering Inc.). Eye fundoscopy was performed before injection and 1, 2, 3, 4- and 6-weeks post-injection to monitor GFP expression over time, using a canon UVI retinal camera connected to a digital imaging system (LHEDIOPH 1600; AMS Ophta).

### 4.12 Whole-mount immunofluorescence in NHP retinas

Eyes of NHPs were enucleated and fixed for 1 hour in 4% paraformaldehyde in phosphate-buffered saline (PBS) solution at RT. Next, the anterior part and the lens was removed and the neuroretina separated from the RPE-choroid-sclera using fine forceps. Retinas were then washed 3 times in PBS at RT and incubated with polyclonal rabbit LM opsin and S opsin antibody (1/300; Vector Laboratories, Peterborough, UK) for 48 hours at 4°C. After 3 washes in PBS, retinas were incubated for 2 hours at RT with 546-conjugated anti-rabbit IgG antibody (1/300; Life Technologies, Saint Aubin, France). Cell nuclei were labelled with Draq5 (1/500; Biostatus, Leicestershire, UK) for 2 hours at RT. Retinas were subsequently washed in PBS, mounted on slides with coverslips in Prolong Gold anti-fade reagent (Life Technologies), and stored at RT before microscopic analysis. Flat-mounted neuroretinas were examined using a laser scanning confocal microscope (Nikon A1RSi). Three-dimensional digital images were collected at 20× (NA 0.75) and 60× (NA 1.4) using NIS-Elements confocal software and appropriate fluorescence filters for laser excitation at 488, 647, and 546 nm.

### 4.13 Immunohistochemistry GFP staining in rats

Rat eyes were enucleated, fixed for 5 hours in Bouin’s solution, dehydrated in successive alcohol solutions and embedded in paraffin. Entire retinas were sliced into 5-μm sections. Paraffin tissue sections were deparaffinized and rehydrated, and antigen was unmasked by boiling the section in ready-to-use citrate-based solution (pH 6.0). Sections were permeabilized for 10 min at RT in PBS containing 0.2% Triton X-100. Non-specific activity was blocked by incubating the sections for 45 minutes at RT in 2% goat serum in PBS and 5% BSA in PBS. Sections were incubated overnight at 4°C in primary antibody diluted in PBS (1:100 monoclonal mouse anti-GFP; 632380, Clontech). After washing with PBS, sections were incubated for 1 h at RT with goat anti-mouse Alexa 555-conjugated antibody (1:300; A21434, Life Technologies). After counterstaining with hematoxylin slides were mounted under coverslips in Prolong Gold anti-fade reagent (Life Technologies). Stained slides were scanned using the Hamamatsu NanoZoomer (Bacus Laboratories, Chicago, IL) and analyzed using NDP.View2 software. Mayer’s hemalum staining images and the corresponding fluorescence image were merged using Fiji software (http://fiji.sc).

### 4.14 Absolute quantification of vector genomes by qPCR in rat tissue samples

Samples (neuroretinas + retinal pigment epithelium (RPE)/choroid) were obtained just after sacrifice in conditions that minimized cross contamination and avoided qPCR inhibition, as described by Le Guiner et al..^28^ Samples were snap-frozen in liquid nitrogen and stored at ≤-70°C before DNA extraction. gDNA was extracted from the whole retinal pigment epithelium using the Nucleospin Tissue Kit (Macherey Nagel) and TissueLyserII (Qiagen), according to the manufacturer’s instructions. qPCR was performed on a StepOne PlusTM Real Time PCR System (Applied Biosystems, Thermo Fisher Scientific) using 50 ng of gDNA in duplicate. Vector genome copy number was determined using the following primer/probe combination, designed to amplify a specific region of the *GFP* sequence present in the transgene (Forward, 5’-ACTACAACAGCCACAACGTCTATATCA-3’; Reverse, 5’-GGCGGATCTTGAAGTTCACC-3’; Probe, 5’-FAM-CCGACAAGCAGAAGAACGGCATCA-TAMRA-3’). Endogenous gDNA copy number was determined using a primer/probe combination designed to amplify the rat *Hprt1* gene (Forward, 5’-GCGAAAGTGGAAAAGCCAAGT -3’; Reverse, 5’-GCCACATCAACAGGACTCTTGTAG -3’; Probe, 5’-FAM-CAAAGCCTAAAAGACAGCGGCAAGTTGAAT -TAMRA-3’). For each sample, cycle threshold (Ct) values were compared with those obtained with different dilutions of linearized standard plasmids (containing either the *eGFP* expression cassette or the rat *Hprt1* gene). The absence of qPCR inhibition in the presence of gDNA was determined by analyzing 50 ng genomic DNA (gDNA) extracted from tissue samples from a control animal and spiked with different dilutions of standard plasmid. Results are expressed as vector genome copy number per diploid genome (vg/dg). The lower limit of quantification (LLOQ) of our test was 0.002 vg/dg. GraphPad Prism 9 software was used for statistical analysis. The nonparametric Mann Whitney test was used.

### 4.15 Relative quantification of GFP expression by RT-qPCR in rat tissue samples

RNA was extracted from neuroretinas and whole RPE, using the Nucleospin RNA kit (Macherey Nagel) and TissueLyserII (Qiagen), according to the manufacturer’s instructions. 1000 ng of total RNA was treated with RNAse-free DNAse I (ezDNAse from Thermo Fisher Scientific) and then reverse transcribed using SuperScript IV Vilo reverse transcriptase (Thermo Fischer Scientific) in a final volume of 20µL. qPCR analysis was then performed on cDNA (diluted 1/20) using the same *GFP* primers and probe as for the quantification of transgene copy number by qPCR. *Hprt1* messenger was also amplified as an endogenous control. For each RNA sample, the absence of DNA contamination was confirmed by analysis of “cDNA-like samples” obtained without adding reverse transcriptase to the reaction mix. The absence of qPCR inhibition in the presence of cDNA was determined by analyzing cDNA obtained from RPE from a control animal, spiked with different dilutions of a standard plasmid. For each RNA sample, Ct values were compared with those obtained with different dilutions of standard plasmids (containing either the *GFP* expression cassette or the *Hprt1* gene). Results were expressed in relative quantities (RQ): RQ = 2-ΔCt = 2-(Ct target - Ct endogenous control). The lower limit of quantification (LLOQ) of our test was RQ = 9^E^-4. GraphPad Prism 9 software was used for statistical analysis. Non-parametric Mann Whitney test was used. Samples were considered significantly different if *p ˂ 0.05, **p ˂ 0.01, ***p ˂ 0.001.

### 4.16 Absolute quantification of vector genomes by qPCR in nonhuman primate tissue samples

Samples (liver, spleen, occipital cortex, and chiasma optic) were obtained just after sacrifice under conditions that minimized cross contamination and avoided qPCR inhibition, as described by Le Guiner et al. (Methods Mol Biol, 2011). Samples were snap-frozen in liquid nitrogen and stored at ≤-70°C before DNA extraction. gDNA was extracted using the Gentra puregene kit and TissueLyserII (both from Qiagen), according to the manufacturer’s instructions. qPCR analyses were conducted on a StepOne PlusTM Real Time PCR System (Applied Biosystems, Thermo Fisher Scientific) using 50 ng of gDNA in duplicate. Vector genome copy number was determined using the same GFP-specific primer/probe combination as used for rat tissue. Endogenous gDNA copy number was determined using a primer/probe combination designed to amplify the macaque *Eglobin* gene (Forward, 5’-TGGCAAGGAGTTCACCCCT -3’; Reverse, 5’-AATGGCGACAGCAGACACC -3’; Probe, 5’-FAM-TGCAGGCTGCCTGGCAGAAGC -TAMRA-3’). For each sample, cycle threshold (Ct) values were compared with those obtained with different dilutions of linearized standard plasmids (containing either the eGFP expression cassette or the macaque *Eglobin* gene). The absence of qPCR inhibition in the presence of gDNA was determined by analyzing 50 ng of gDNA extracted from tissue samples from a control animal and spiked with different dilutions of standard plasmid. Results are expressed as vector genome copy number per diploid genome (vg/dg). The lower limit of quantification (LLOQ) of our test was 0.001 vg/dg.

## Supporting information

complete supplemental file

## Acknowledgements

The authors thank all personnel at the Boisbonne Center for Gene Therapy (ONIRIS, INSERM, Nantes, France) for handling and care of the rats and nonhuman primates included in this study. We also thank the vector core of TarGeT, UMR 1089 (CPV, INSERM and Nantes Université, http://umr1089.univ-nantes.fr) for the production of the rAAV vectors used in this study, and the staff at the Preclinical Analytics Core of TarGeT, UMR 1089 (PAC, INSERM and Nantes Université) for the molecular analyses performed in this study. Thanks also to the IBISA MicroPICell facility (Biogenouest), a member of the France-Bioimaging national infrastructure supported by the French national research agency (ANR-10-INBS-04). Analysis and purification of chemical compounds were performed at the Chromatography Platform, CEISAM laboratory, UMR 6230 CNRS/UN, Nantes Université. Thanks to Nicolas Ferry for reviewing the manuscript and to Owen Howard PhD for English language editing. This research was supported by the Fondation d’Entreprise Thérapie Génique en Pays de Loire, the Centre Hospitalier Universitaire (CHU) of Nantes, the Institut National de la Santé et de la Recherche Médicale (INSERM), and Nantes Université, by a grant from the French National Agency for Research (“Investissements d’Avenir” Equipex ArronaxPlus n°ANR-11-EQPX-0004) and by Coave Therapeutics (formerly Horama).

## Conflicts of Interest

M.M., D.D, E.A. are inventors on a patent including the technology described in this manuscript. AG, NB and GML are employees of Coave Therapeutics.

The authors have no potential conflicts of interest to disclose.

## Data availability

All relevant data are available from the authors on request and are included in the manuscript.

## Appendix A. Supplementary data

^1^H NMR, ^13^C NMR, analytical HPLC spectra, bioconjugation characterization, in vivo retina analyses.

## Notes

### Competing Interest Statement

The authors have declared no competing interest.

## References

1. Prado, D.A., Acosta-Acero, M., and Maldonado, R.S. (2020). Gene therapy beyond luxturna: a new horizon of the treatment for inherited retinal disease. Curr Opin Ophthalmol 31, 147–154. 10.1097/Icu.0000000000000660.

2. Voretigene Neparvovec-rzyl (Luxturna) for Inherited Retinal Dystrophy. (2018). Med Lett Drugs Ther 60, 53–55.

3. Hudson, N., and Campbell, M. (2019). Inner Blood-Retinal Barrier Regulation in Retinopathies. Advances in experimental medicine and biology 1185, 329–333. 10.1007/978-3-030-27378-1_54.

4. Samiy, N. (2014). Gene therapy for retinal diseases. Journal of ophthalmic & vision research 9, 506–509. 10.4103/2008-322X.150831.

5. Niederkorn, J.Y. (2019). The Eye Sees Eye to Eye With the Immune System: The 2019 Proctor Lecture. Investigative ophthalmology & visual science 60, 4489–4495. 10.1167/iovs.19-28632.

6. Chan, Y.K., Dick, A.D., Hall, S.M., Langmann, T., Scribner, C.L., Mansfield, B.C., and w, O.G.T.I. (2021). Inflammation in Viral Vector-Mediated Ocular Gene Therapy: A Review and Report From a Workshop Hosted by the Foundation Fighting Blindness, 9/2020. Transl Vis Sci Techn 10. 10.1167/tvst.10.4.3.

7. Petrs-Silva, H., Dinculescu, A., Li, Q., Min, S.H., Chiodo, V., Pang, J.J., Zhong, L., Zolotukhin, S., Srivastava, A., Lewin, A.S., and Hauswirth, W.W. (2009). High-efficiency transduction of the mouse retina by tyrosine-mutant AAV serotype vectors. Mol Ther 17, 463–471. 10.1038/mt.2008.269.

8. Petrs-Silva, H., Dinculescu, A., Li, Q., Deng, W.T., Pang, J.J., Min, S.H., Chiodo, V., Neeley, A.W., Govindasamy, L., Bennett, A., Agbandje-McKenna, M., et al. (2011). Novel properties of tyrosine-mutant AAV2 vectors in the mouse retina. Mol Ther 19, 293–301. 10.1038/mt.2010.234.

9. Dalkara, D., Byrne, L.C., Klimczak, R.R., Visel, M., Yin, L., Merigan, W.H., Flannery, J.G., and Schaffer, D.V. (2013). In vivo-directed evolution of a new adeno-associated virus for therapeutic outer retinal gene delivery from the vitreous. Science translational medicine 5, 189ra176. 10.1126/scitranslmed.3005708.

10. Buning, H., and Srivastava, A. (2019). Capsid Modifications for Targeting and Improving the Efficacy of AAV Vectors. Molecular therapy. Methods & clinical development 12, 248–265. 10.1016/j.omtm.2019.01.008.

11. Khabou, H., Desrosiers, M., Winckler, C., Fouquet, S., Auregan, G., Bemelmans, A.P., Sahel, J.A., and Dalkara, D. (2016). Insight into the mechanisms of enhanced retinal transduction by the engineered AAV2 capsid variant-7m8. Biotechnology and Bioengineering 113, 2712–2724. 10.1002/bit.26031.

12. Mevel, M., Bouzelha, M., Leray, A., Pacouret, S., Guilbaud, M., Penaud-Budloo, M., Alvarez-Dorta, D., Dubreil, L., Gouin, S.G., Combal, J.P., Hommel, M., et al. (2019). Chemical modification of the adeno-associated virus capsid to improve gene delivery. Chemical science 11, 1122–1131. 10.1039/c9sc04189c.

13. Free, P., Hurley, C.A., Kageyama, T., Chain, B.M., and Tabor, A.B. (2006). Mannose-pepstatin conjugates as targeted inhibitors of antigen processing. Org Biomol Chem 4, 1817–1830. 10.1039/b600060f.

14. Boye, S.E., Alexander, J.J., Witherspoon, C.D., Boye, S.L., Peterson, J.J., Clark, M.E., Sandefer, K.J., Girkin, C.A., Hauswirth, W.W., and Gamlin, P.D. (2016). Highly Efficient Delivery of Adeno-Associated Viral Vectors to the Primate Retina. Hum Gene Ther 27, 580–597. 10.1089/hum.2016.085.

15. Sugawara, K., Hirabayashi, G., Kamiya, N., and Kuramitz, H. (2006). Evaluation of concanavalin A-mannose interaction on the electrode covered with collagen film. Talanta 68, 1176–1181. 10.1016/j.talanta.2005.07.036.

16. Wright, J.F., Le, T., Prado, J., Bahr-Davidson, J., Smith, P.H., Zhen, Z., Sommer, J.M., Pierce, G.F., and Qu, G. (2005). Identification of factors that contribute to recombinant AAV2 particle aggregation and methods to prevent its occurrence during vector purification and formulation. Mol Ther 12, 171–178. 10.1016/j.ymthe.2005.02.021.

17. Vandenberghe, L.H., Bell, P., Maguire, A.M., Cearley, C.N., Xiao, R., Calcedo, R., Wang, L., Castle, M.J., Maguire, A.C., Grant, R., Wolfe, J.H., et al. (2011). Dosage thresholds for AAV2 and AAV8 photoreceptor gene therapy in monkey. Science translational medicine 3, 88ra54. 10.1126/scitranslmed.3002103.

18. Maguire, A.M., High, K.A., Auricchio, A., Wright, J.F., Pierce, E.A., Testa, F., Mingozzi, F., Bennicelli, J.L., Ying, G.S., Rossi, S., Fulton, A., et al. (2009). Age-dependent effects of RPE65 gene therapy for Leber’s congenital amaurosis: a phase 1 dose-escalation trial. Lancet 374, 1597–1605. 10.1016/S0140-6736(09)61836-5.

19. Wiley, L.A., Burnight, E.R., Kaalberg, E.E., Jiao, C., Riker, M.J., Halder, J.A., Luse, M.A., Han, I.C., Russell, S.R., Sohn, E.H., Stone, E.M., et al. (2018). Assessment of Adeno-Associated Virus Serotype Tropism in Human Retinal Explants. Hum Gene Ther 29, 424–436. 10.1089/hum.2017.179.

20. Han, I.C., Cheng, J.L., Burnight, E.R., Ralston, C.L., Fick, J.L., Thomsen, G.J., Tovar, E.F., Russell, S.R., Sohn, E.H., Mullins, R.F., Stone, E.M., et al. (2020). Retinal Tropism and Transduction of Adeno-Associated Virus Varies by Serotype and Route of Delivery (Intravitreal, Subretinal, or Suprachoroidal) in Rats. Hum Gene Ther 31, 1288–1299. 10.1089/hum.2020.043.

21. Picaud, S., Dalkara, D., Marazova, K., Goureau, O., Roska, B., and Sahel, J.A. (2019). The primate model for understanding and restoring vision. Proc Natl Acad Sci U S A. 10.1073/pnas.1902292116.

22. Russell, S., Bennett, J., Wellman, J.A., Chung, D.C., Yu, Z.F., Tillman, A., Wittes, J., Pappas, J., Elci, O., McCague, S., Cross, D., et al. (2017). Efficacy and safety of voretigene neparvovec (AAV2-hRPE65v2) in patients with RPE65-mediated inherited retinal dystrophy: a randomised, controlled, open-label, phase 3 trial. Lancet 390, 849–860. 10.1016/S0140-6736(17)31868-8.

23. Weed, L., Ammar, M.J., Zhou, S., Wei, Z., Serrano, L.W., Sun, J., Lee, V., Maguire, A.M., Bennett, J., and Aleman, T.S. (2019). Safety of Same-Eye Subretinal Sequential Readministration of AAV2-hRPE65v2 in Non-human Primates. Molecular therapy. Methods & clinical development 15, 133–148. 10.1016/j.omtm.2019.08.011.

24. MacLaren, R.E., Groppe, M., Barnard, A.R., Cottriall, C.L., Tolmachova, T., Seymour, L., Clark, K.R., During, M.J., Cremers, F.P., Black, G.C., Lotery, A.J., et al. (2014). Retinal gene therapy in patients with choroideremia: initial findings from a phase 1/2 clinical trial. Lancet 383, 1129–1137. 10.1016/S0140-6736(13)62117-0.

25. Provost, N., Le Meur, G., Weber, M., Mendes-Madeira, A., Podevin, G., Cherel, Y., Colle, M.-A., Deschamps, J.-Y., Moullier, P., and Rolling, F. (2005). Biodistribution of rAAV Vectors Following Intraocular Administration: Evidence for the Presence and Persistence of Vector DNA in the Optic Nerve and in the Brain. Molecular Therapy 11, 275–283. 10.1016/j.ymthe.2004.09.022.

26. Seitz, I.P., Michalakis, S., Wilhelm, B., Reichel, F.F., Ochakovski, G.A., Zrenner, E., Ueffing, M., Biel, M., Wissinger, B., Bartz-Schmidt, K.U., Peters, T., et al. (2017). Superior Retinal Gene Transfer and Biodistribution Profile of Subretinal Versus Intravitreal Delivery of AAV8 in Nonhuman Primates. Investigative Opthalmology & Visual Science 58. 10.1167/iovs.17-22473.

27. D’Costa, S., Blouin, V., Broucque, F., Penaud-Budloo, M., Francois, A., Perez, I.C., Le Bec, C., Moullier, P., Snyder, R.O., and Ayuso, E. (2016). Practical utilization of recombinant AAV vector reference standards: focus on vector genomes titration by free ITR qPCR. Mol Ther Methods Clin Dev 5, 16019. 10.1038/mtm.2016.19.

28. Guiner, C.L., Moullier, P., and Arruda, V.R. (2012). Biodistribution and Shedding of AAV Vectors. In Adeno-Associated Virus, pp. 339–359. 10.1007/978-1-61779-370-7_15.

