## Supplementary material for "Mannose-coupled AAV2: a second generation AAV vector for increased retinal gene therapy efficiency": complete supplemental file

Electronic Supplementary Information (ESI) for

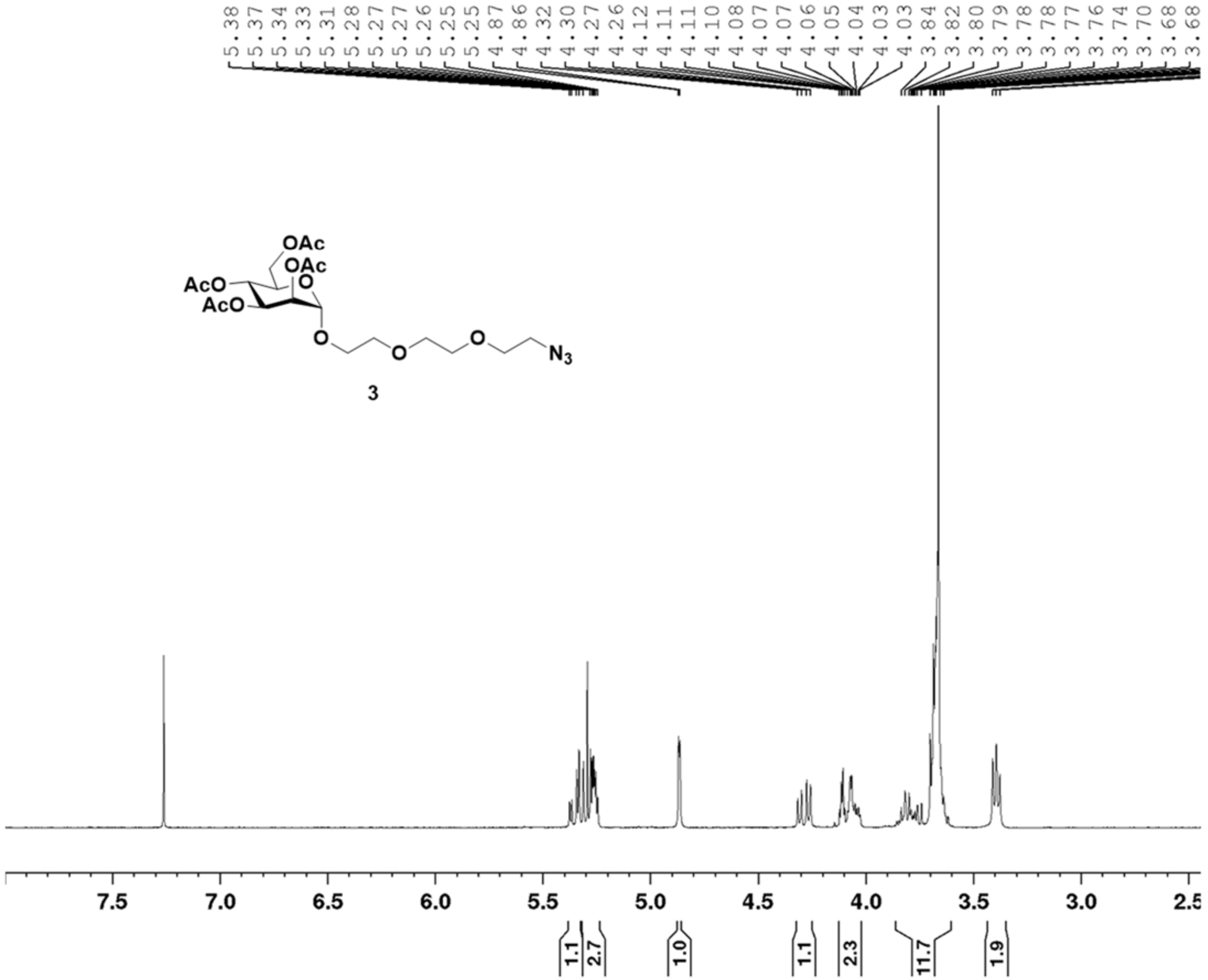

**Fig. S1**: $^{1}$H NMR of compound **3.**

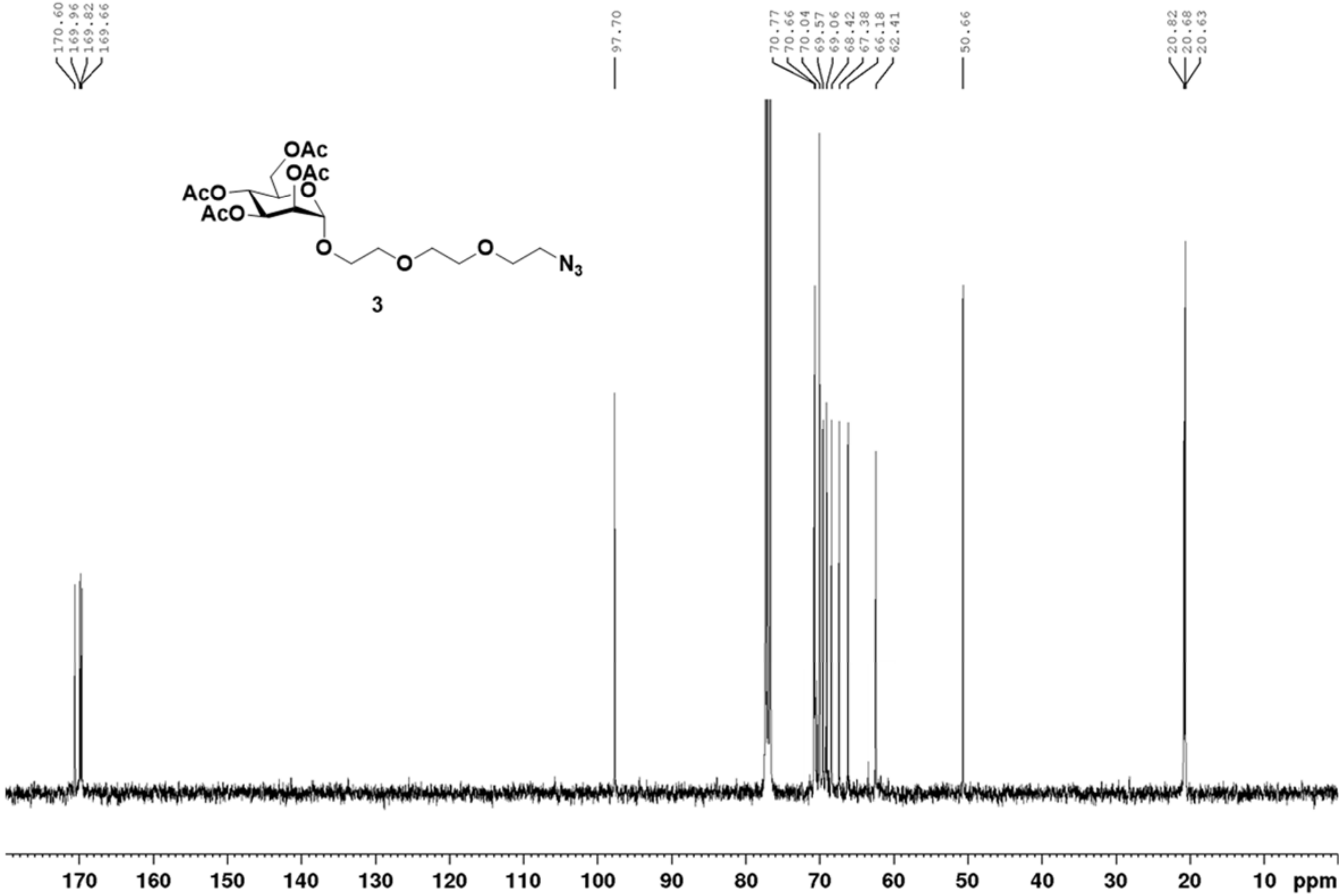


**Fig. S2**: $^{13}$C NMR of compound **3.**

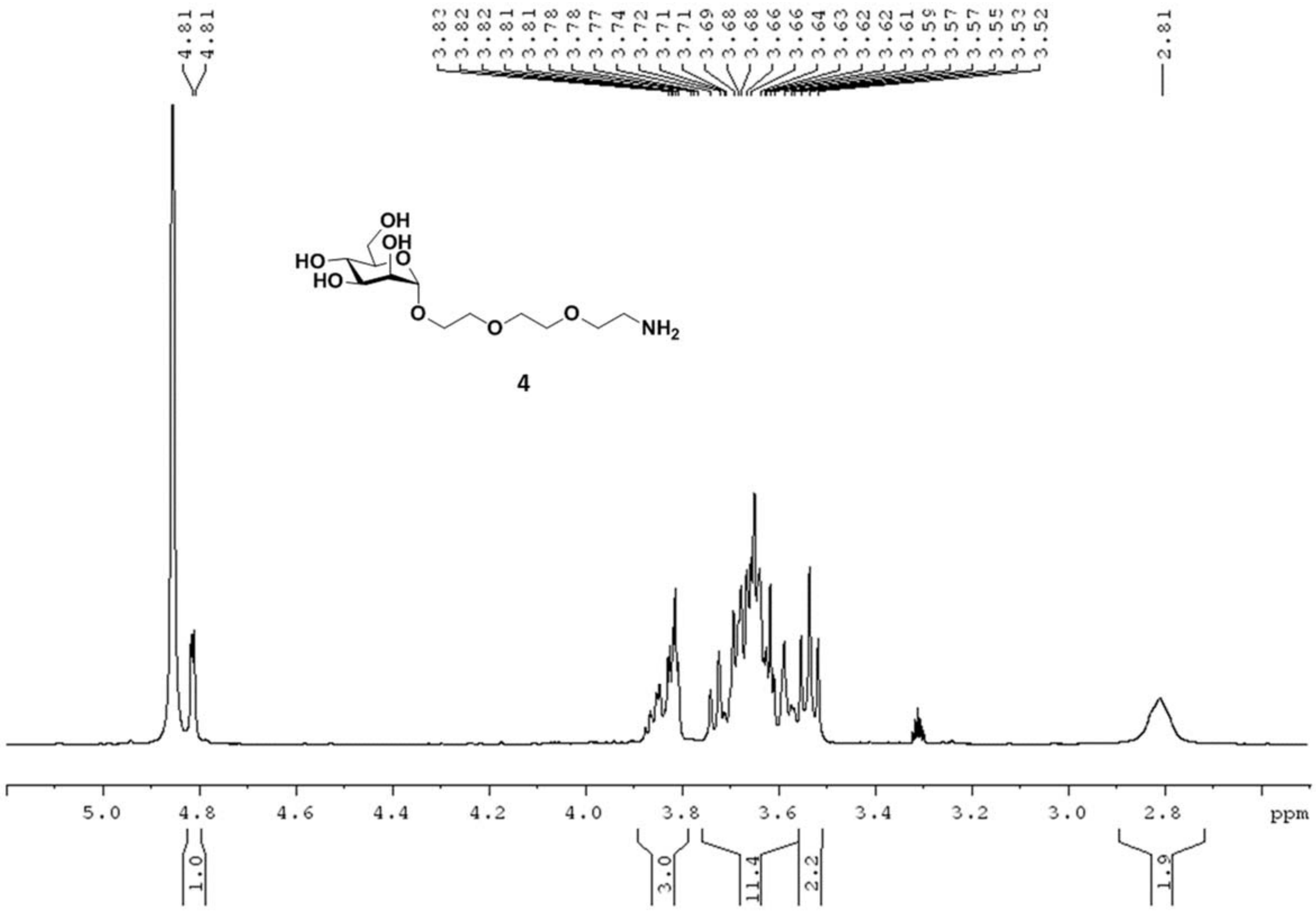


**Fig. S3**: $^1$H NMR of compound **4.**

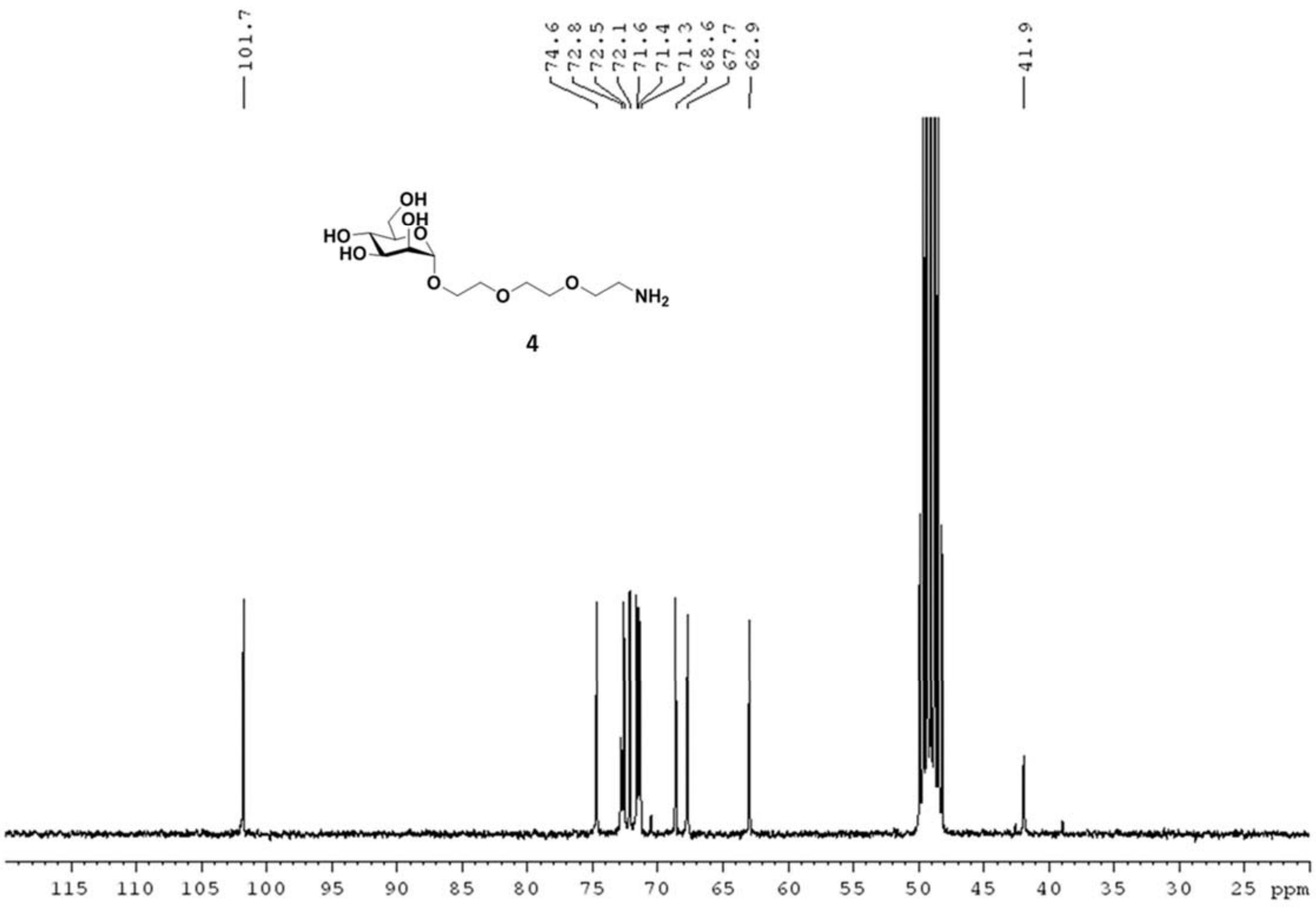


**Fig. S4**: $^{13}C$ NMR of compound **4.**

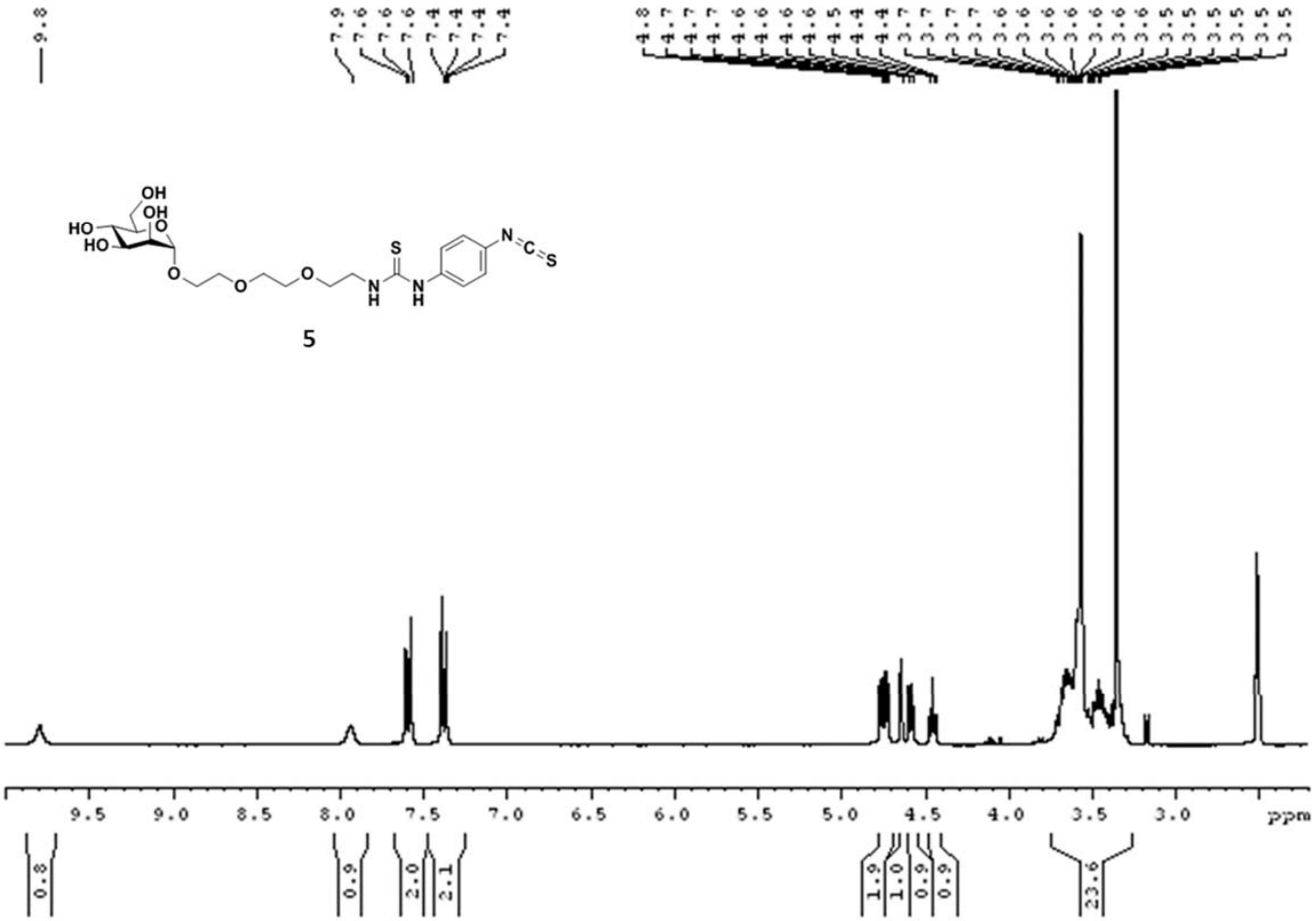


**Fig. S5**: $^{1}$H NMR of compound **5.**

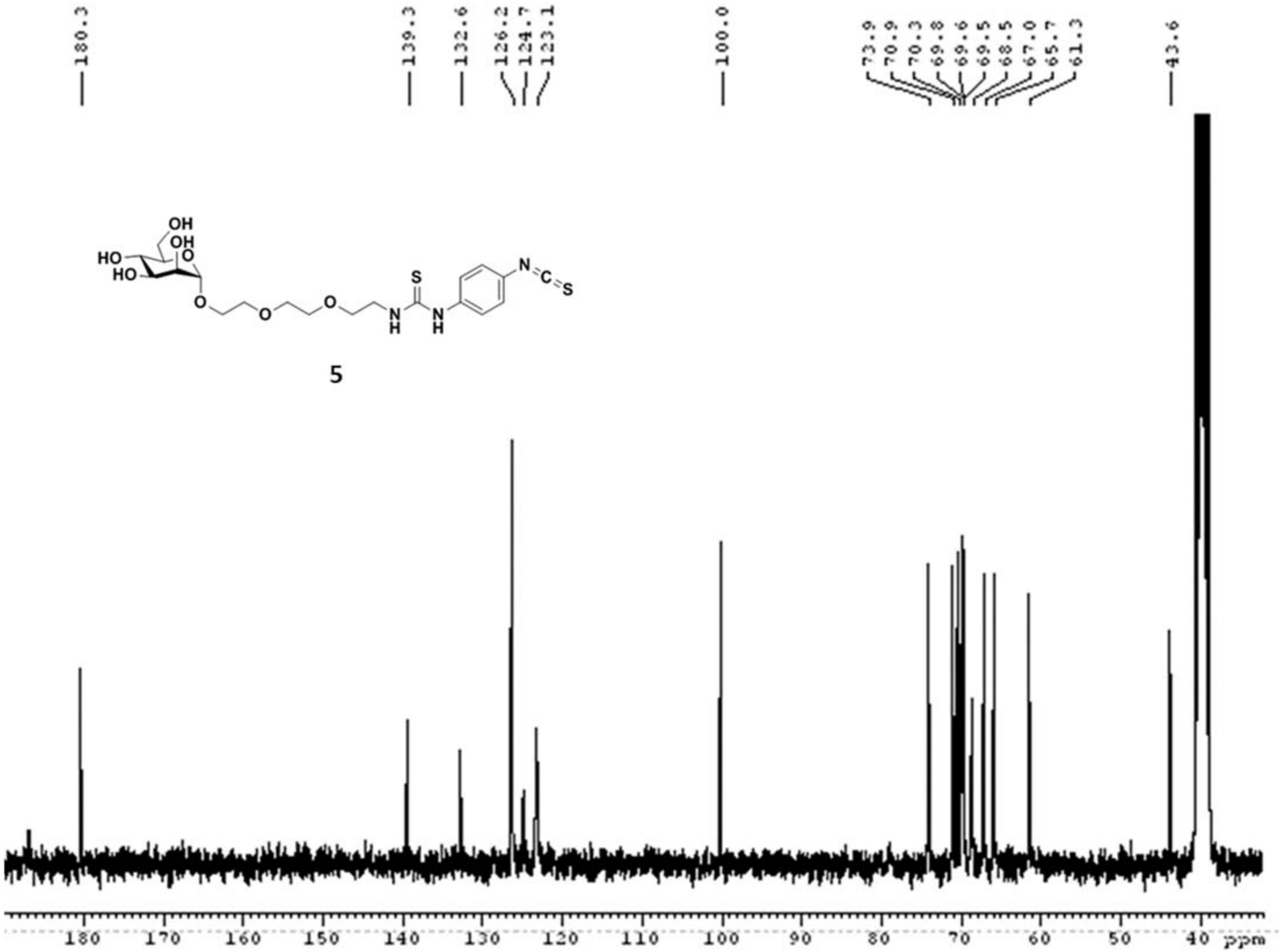


**Fig. S6**: $^{13}$C NMR of compound **5.**

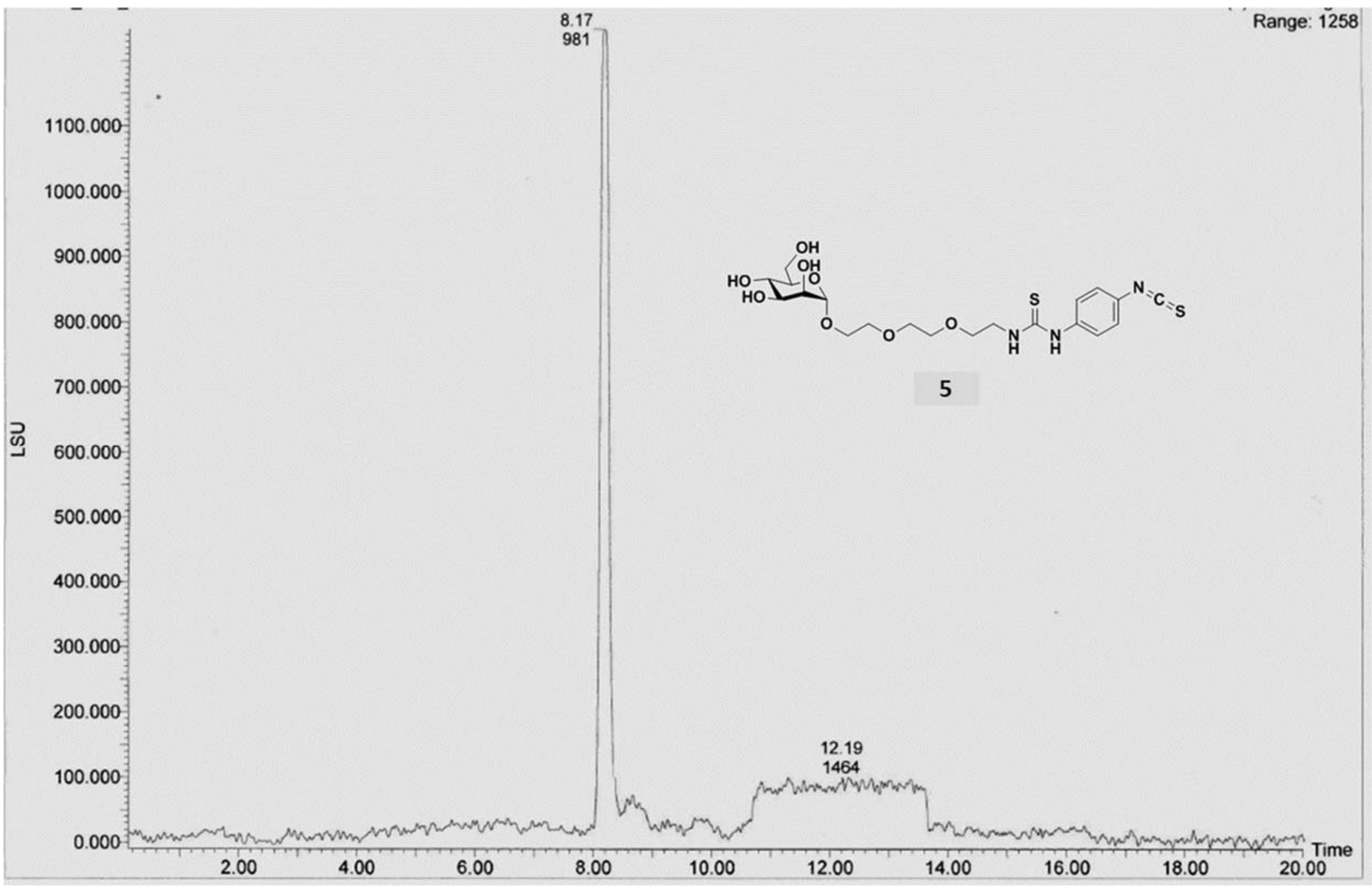


**Fig. S7**: Analytical HPLC of compound **5.**

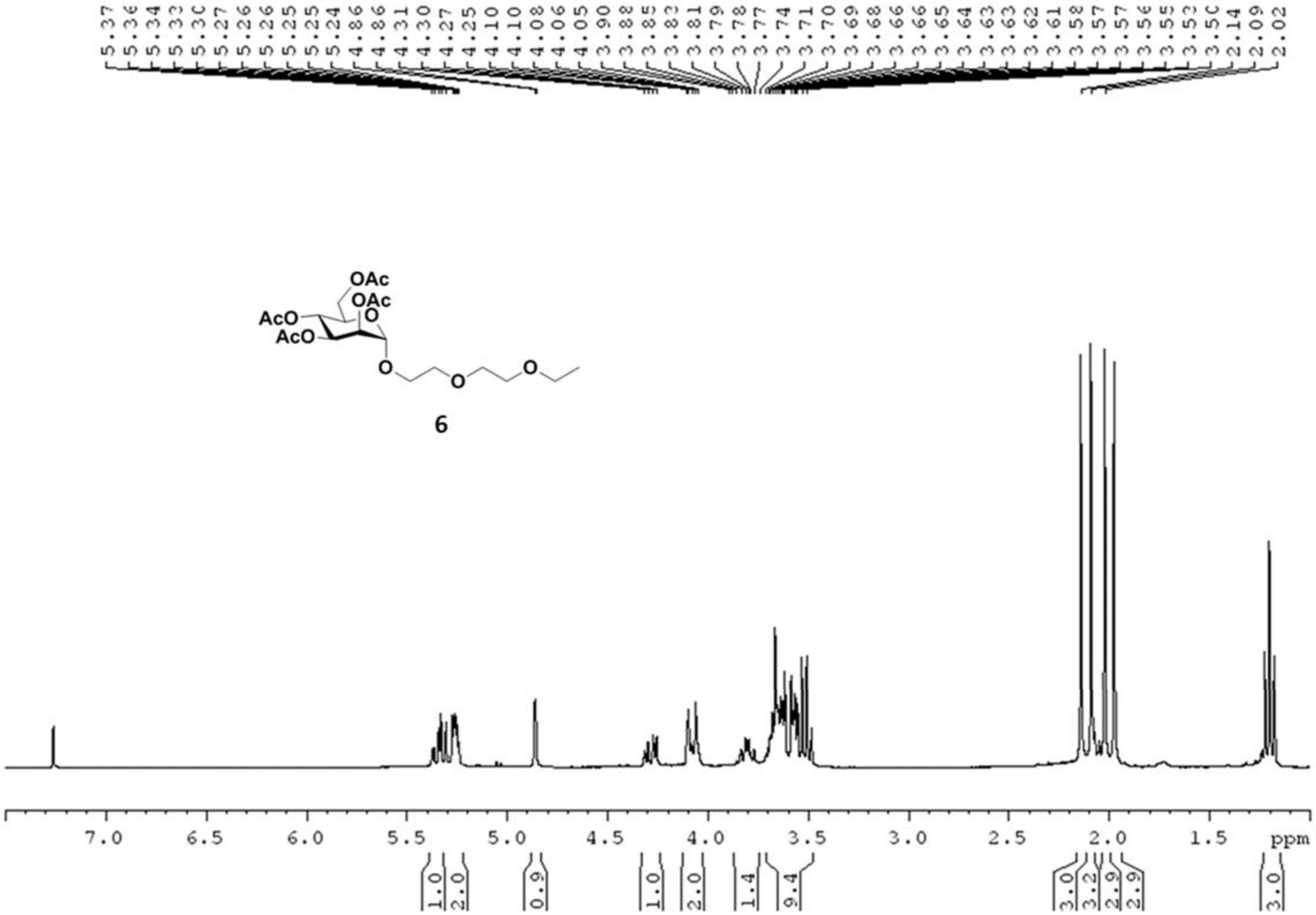


**Fig. S8**: [1]H NMR of compound **6.**

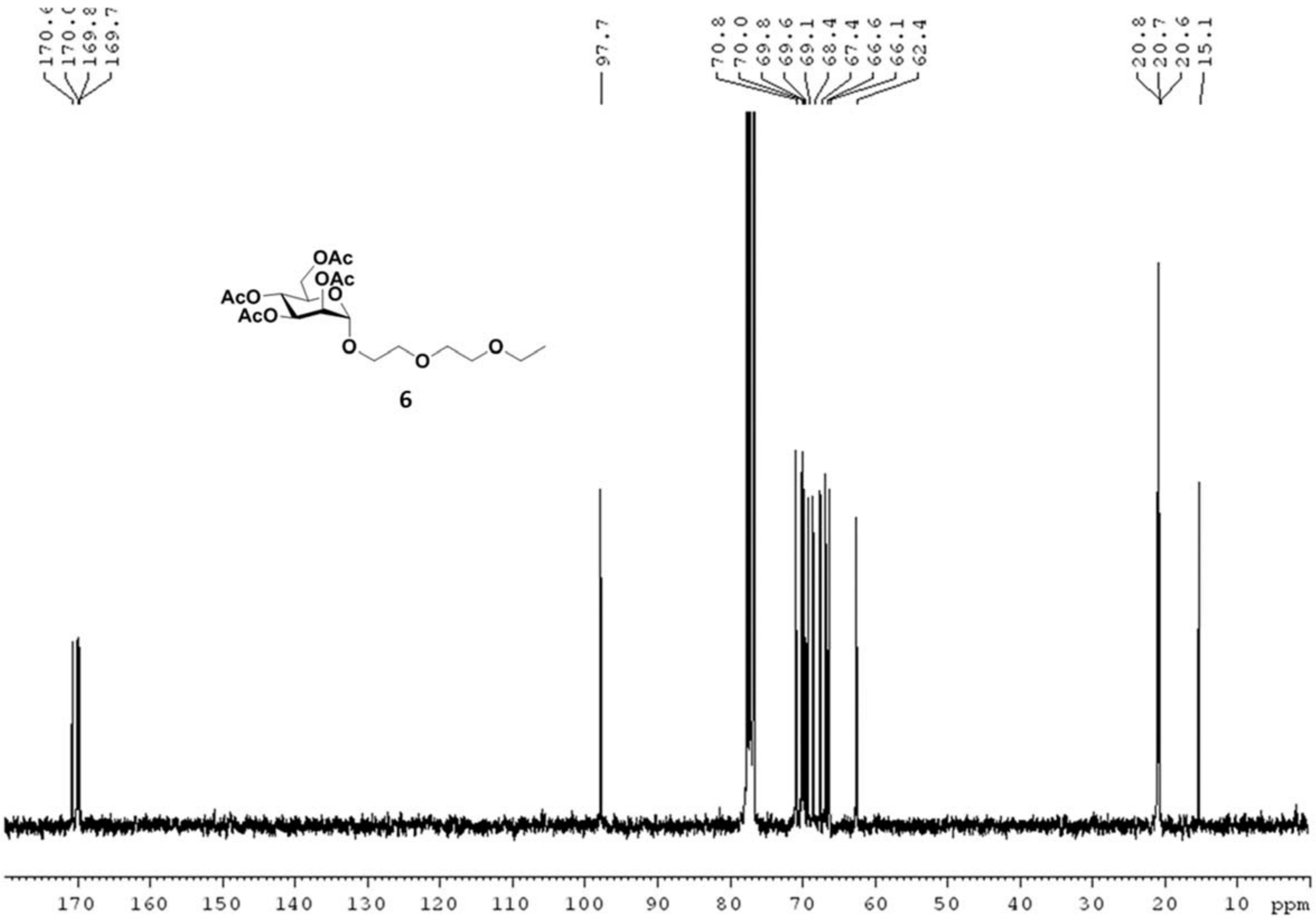


**Fig. S9**: $^{13}$C NMR of compound **6.**

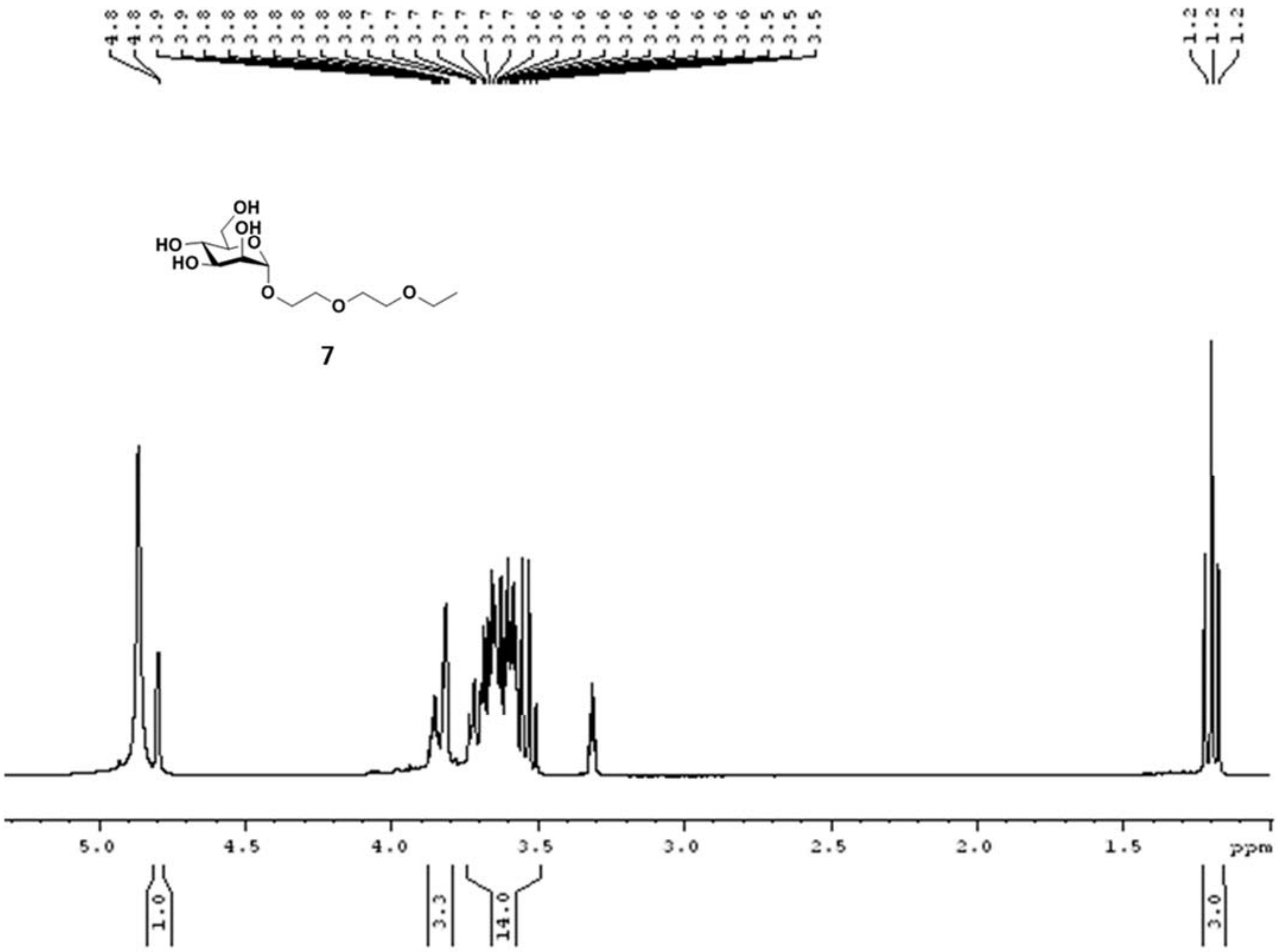


**Fig. S10**: $^{1}$H NMR of compound **7.**

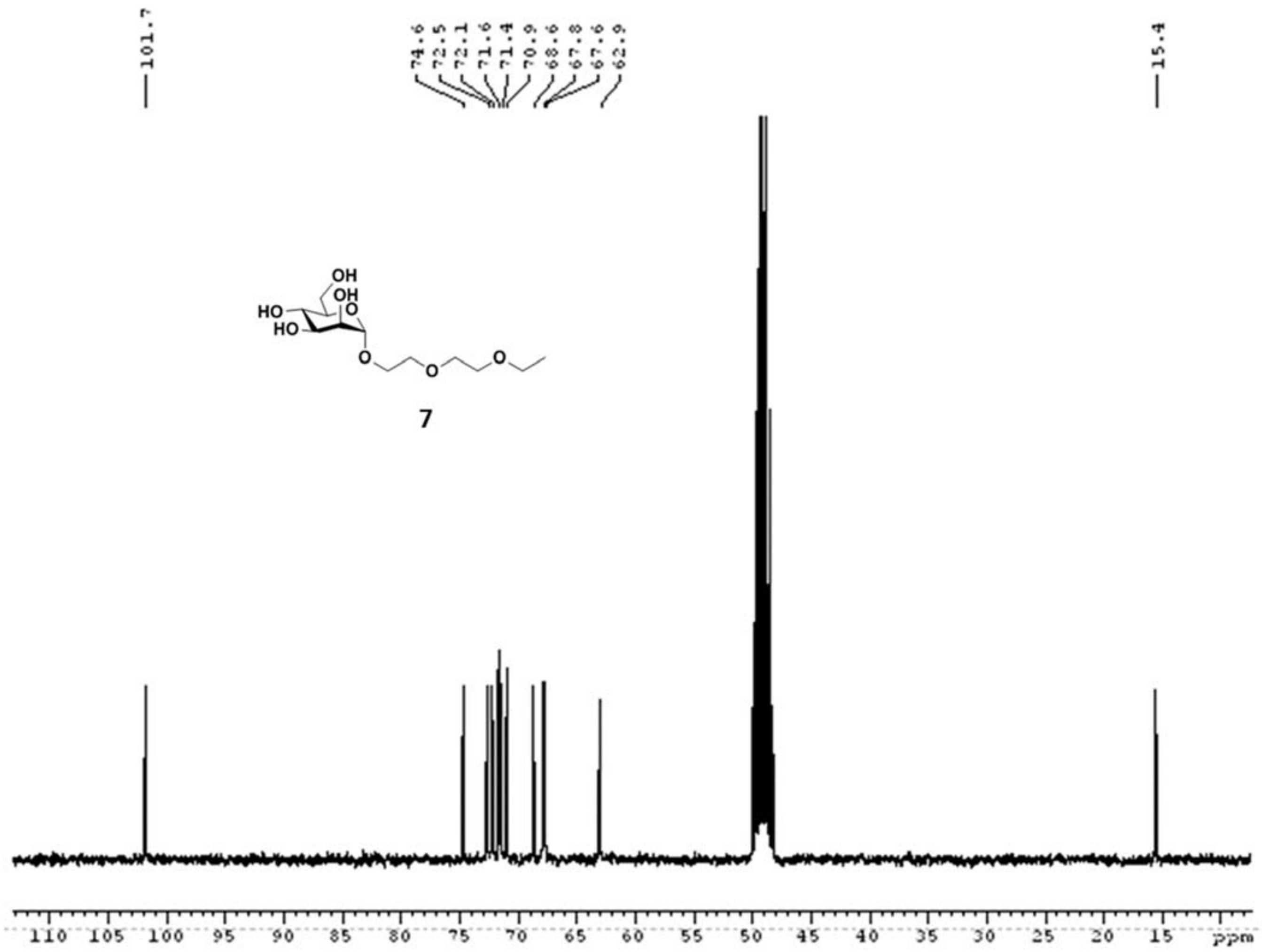


**Fig. S11**: $^{13}$C NMR of compound **7.**

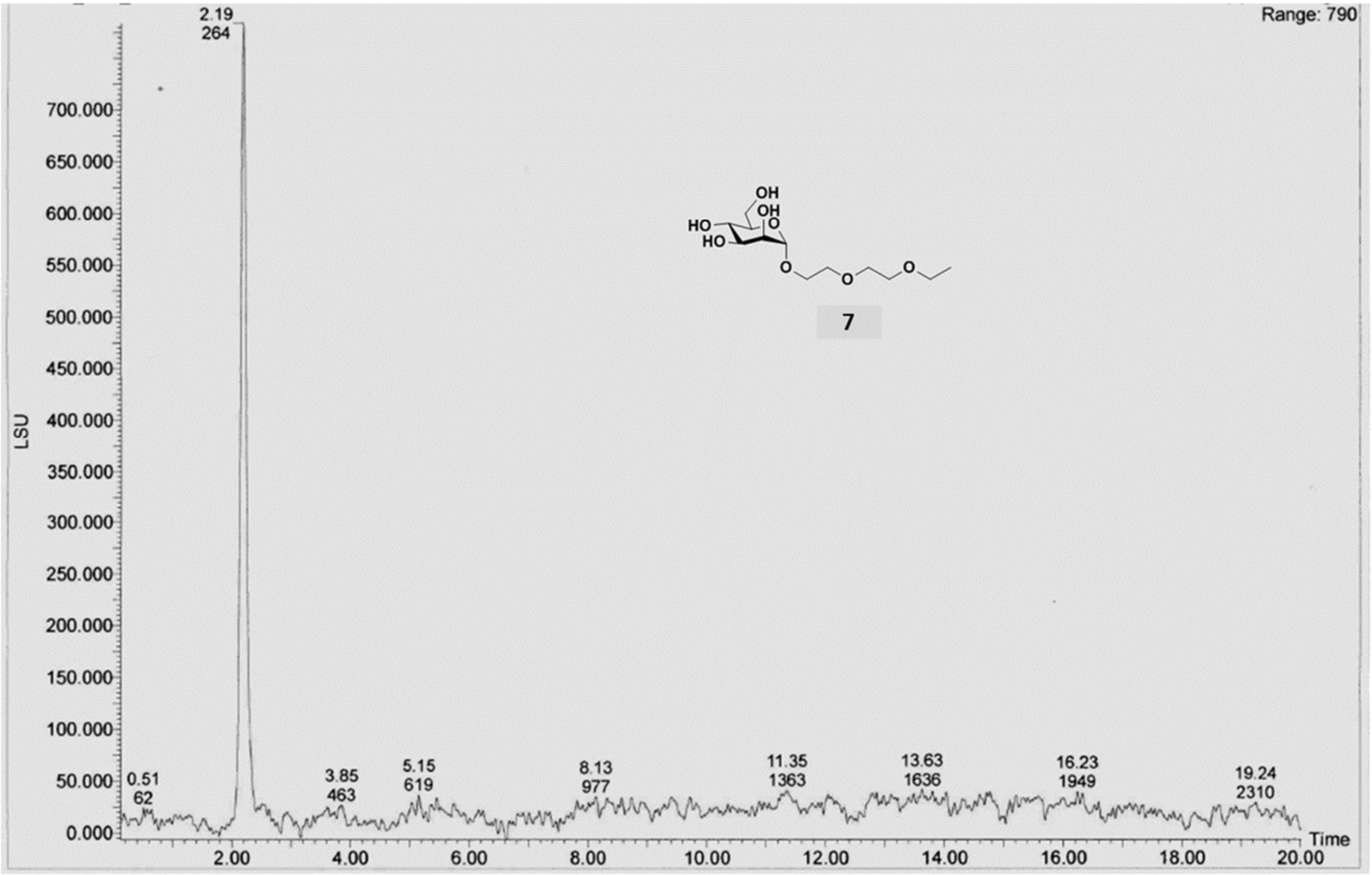


**Fig. S12**: Analytical HPLC of compound **7.**

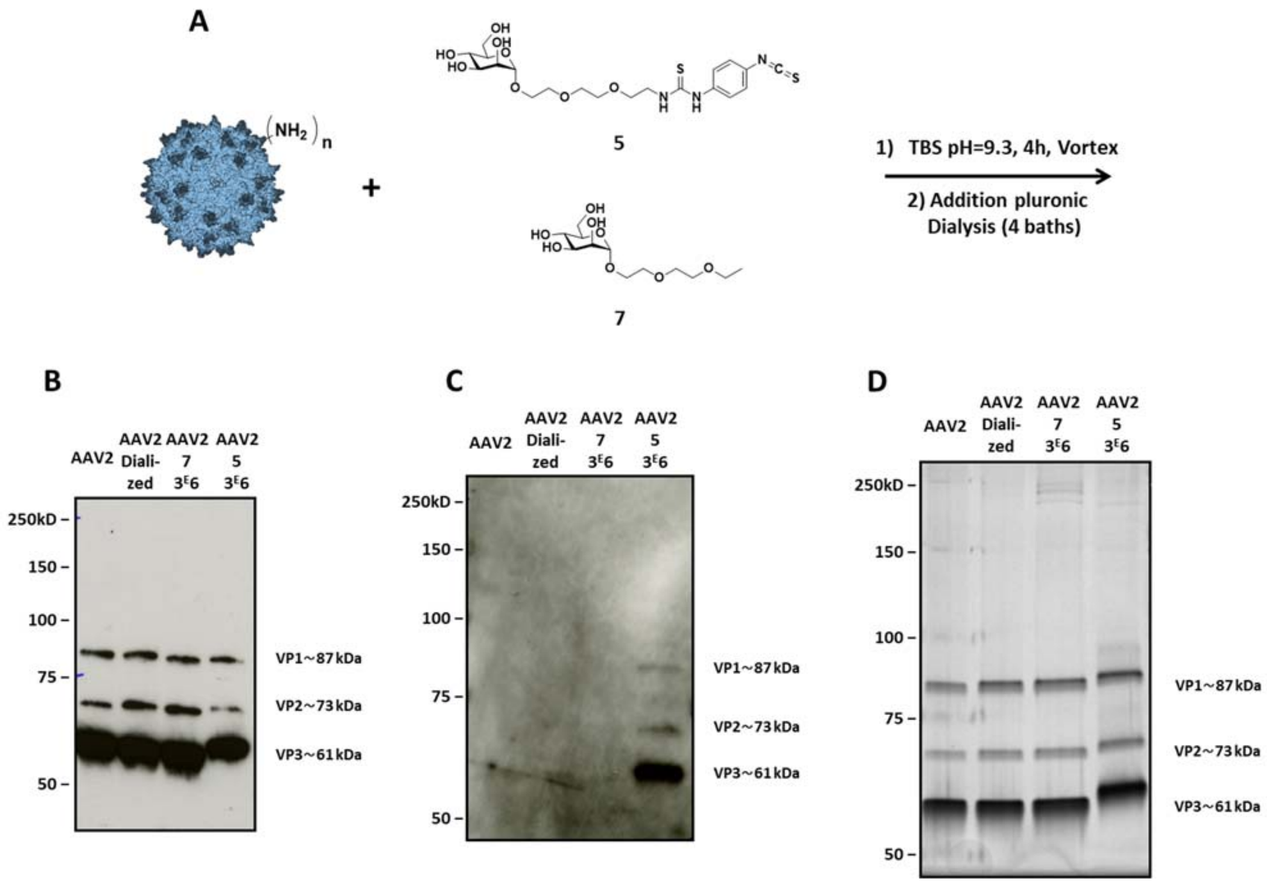


**Fig. S13: Second covalent coupling of compound 5 to the AAV2 capsid *via* primary amino groups.** (**A**) $1^{E}12$ vg of AAV2-GFP vectors were added to a solution of compound **5** ($3^{E}5$ or $3^{E}6$ eq) in TBS buffer (pH 9.3) and incubated for 4 h at RT. The same experimental procedure was followed with compound **7** ($3^{E}6$ eq) in TBS at pH 9.3 as a control. (**B, C**). $5^{E}8$ vg of the samples were analyzed by Western blot using a polyclonal antibody against the capsid to detect VP proteins **(B)** or using FITC-concanavalin A lectin **(C).** Note that compound **7** is a negative control lacking the Ar-NCS reactive function. (**D**) $1^{E}10$ vg of each condition were analyzed by silver nitrate staining. VP1, VP2 and VP3 are the three proteins constituting the AAV capsid. Capsid protein molecular weights are indicated on the right-hand side of the images.

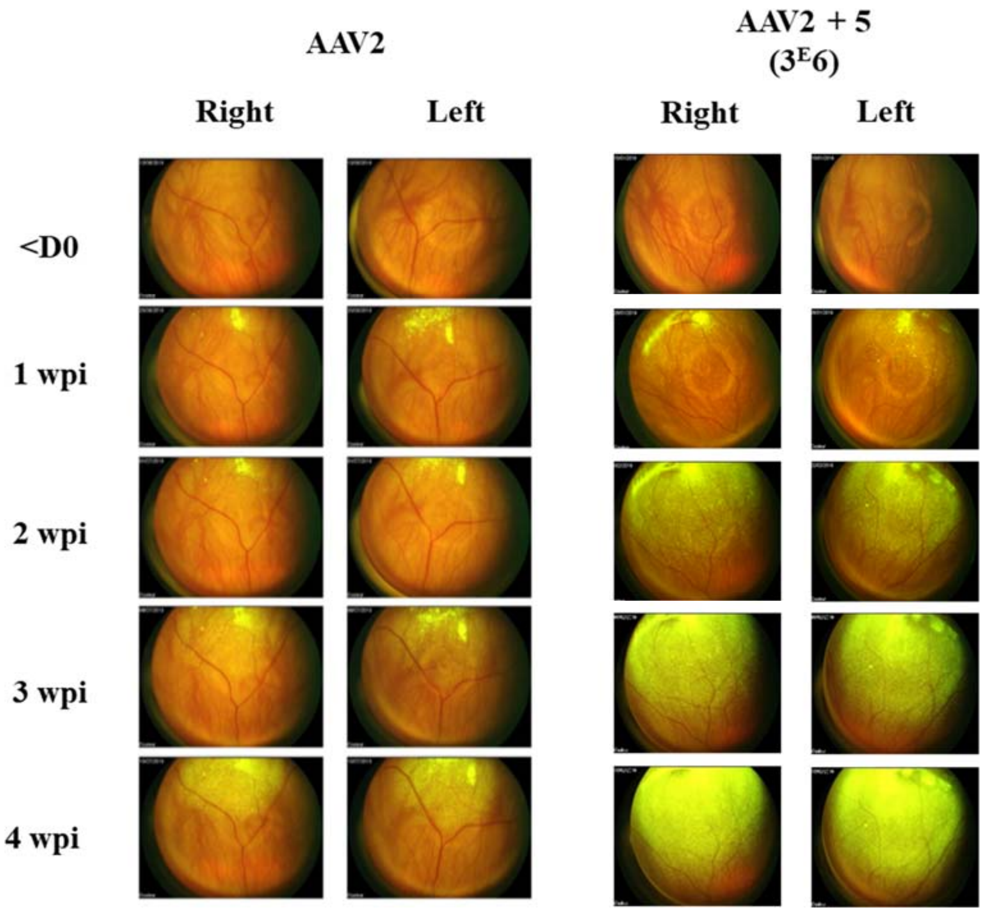


**Fig. S14: Ocular fundus in rats (second coupling).** *In vivo* imaging shows the kinetics of GFP expression after subretinal delivery of AAV2 or AAV2 + **5** ($3^E6$). Representative images obtained in one animal injected with each type of vector are shown. Wpi, weeks post-injection.

**A.**

**B.**

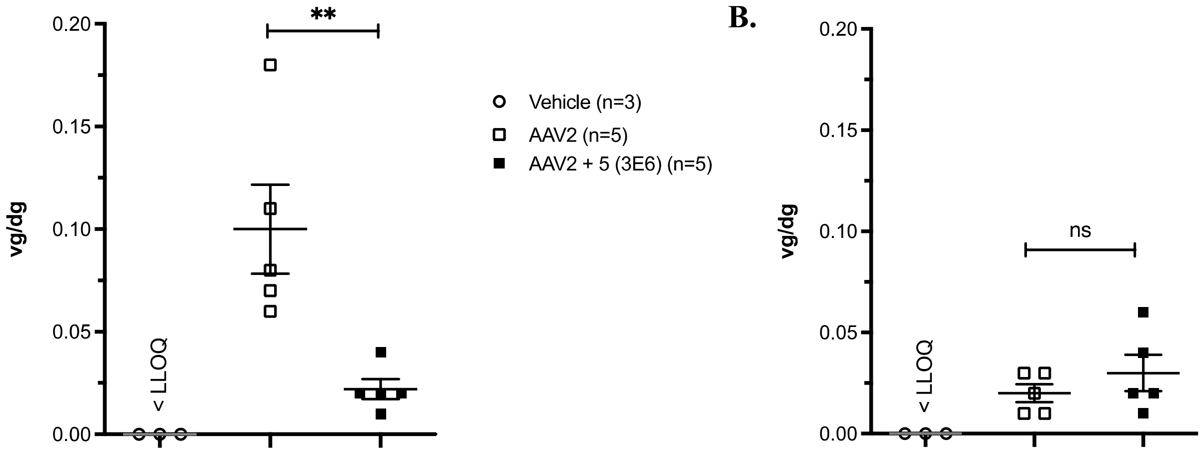


**Fig. S15:** Vector genome copy number per diploid genome (vg/dg) in right neuroretina (A) and RPE (B) measured 4 weeks post injection in rats injected with vehicle, AAV2, or AAV2 + **5** ($3^{E}6$) (second coupling). Results are expressed as the mean ± SEM. The Mann-Whitney U-test was used for statistical analyses: ** $p<0.01$; ns = not significant. Lower limit of quantification (LLOQ) = 0.002 vg/dg.

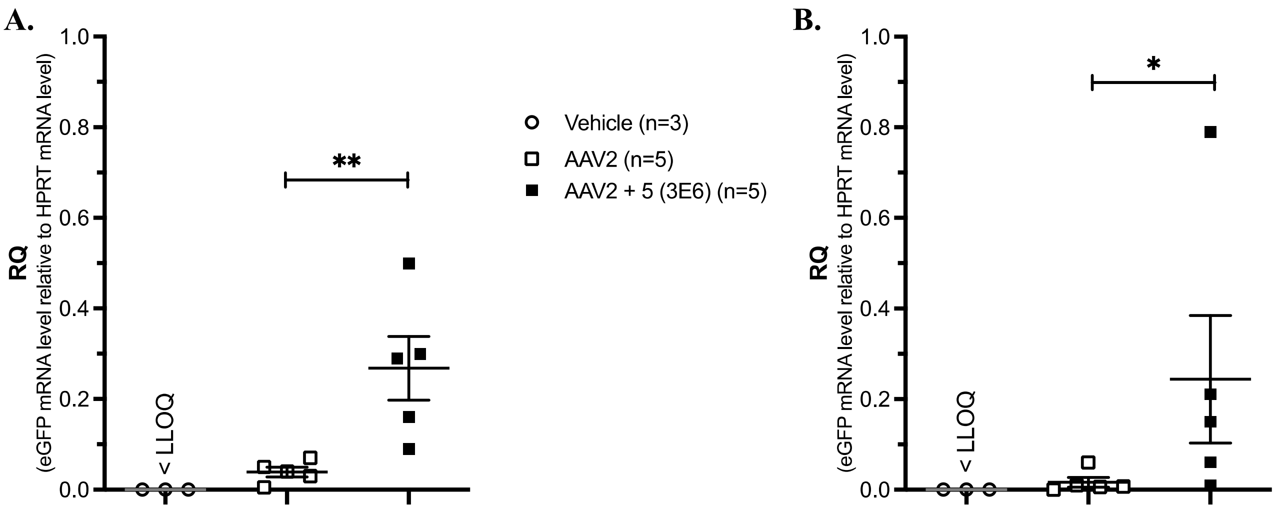


**Fig. S16:** Relative quantification of *GFP* mRNA levels (relative to rat *Hprt1* mRNA levels) analysed by RTqPCR in the left neuroretina (A) and RPE (B) 4 weeks post injection in rats injected with vehicle, AAV2, or AAV2 + **5** ($3^{E}6$) (second coupling). Results are expressed as mean ± SEM. The Mann-Whitney U-test was used for statistical analyses: **p<0.01. Lower limit of quantification (LLOQ) = $3^{E}$-4.

| Animals | Vector | Occulomotor muscles | Optic chiasm | Occipital cortex | Liver | Spleen |
|---|---|---|---|---|---|---|
| NHP 1 | Vehicle | < 0.001 | < 0.001 | < 0.001 | < 0.001 | < 0.001 |
| NHP 2 | AAV2 | < 0.001 | < 0.001 | < 0.001 | < 0.001 | < 0.001 |
| NHP 3 | AAV2 + **5** ($3^E6$) | < 0.001 | **0.002** | < 0.001 | < 0.001 | < 0.001 |

**Table S1:** Vector genome copy number per diploid genome (vg/dg) in tissues from nonhuman primates 4 weeks after subretinal delivery of vehicle, AAV2, or AAV2+**5** ($3^E6$). Lower limit of quantification (LLOQ) = 0.001 vg/dg.
